# Distinct Subsets of Multi-Lymphoid Progenitors Support Ontogeny-Related Changes in Human Lymphopoiesis

**DOI:** 10.1101/2022.12.12.520060

**Authors:** Seydou Keita, Samuel Diop, Shalva Lekiashvili, Emna Chabaane, Elisabeth Nelson, Marion Strullu, Chloé Arfeuille, Fabien Guimiot, Thomas Domet, Sophie Duchez, Bertrand Evrard, Thomas Darde, Jerome Larghero, Els Verhoeyen, Ana Cumano, Elizabeth A. Macintyre, François Jouen, Michele Goodhardt, David Garrick, Frederic Chalmel, Kutaiba Alhaj Hussen, Bruno Canque

## Abstract

Changes in lymphocyte production patterns occurring across human ontogeny remain poorly defined. In this study, we demonstrate that human lymphopoiesis is supported by three waves of embryonic, fetal, and postnatal multi-lymphoid progenitors (MLPs) differing in CD7 and CD10 expression and their output of CD127^-/+^ early lymphoid progenitors (ELP). Our results reveal that, like the fetal-to-adult switch in erythropoiesis, transition to postnatal life coincides with a shift from multilineage to B lineage-biased lymphopoiesis and an increase in production of CD127^+^ ELPs which persists until puberty. A further developmental transition is observed in elderly individuals where B-cell differentiation bypasses the CD127^+^ compartment and branches directly from CD10^+^ MLPs. Functional analyses indicate that these changes are determined at the level of the hematopoietic stem cell. Besides reconciling controversies about the identity and function of human MLPs, these results may shed light on the causes of age-related differences in the incidence of lymphoblastic leukemia.

## INTRODUCTION

The concept of developmental stratification of the immune system was first proposed by Herzenberg & Herzenberg based on studies demonstrating the fetal origin of murine B1 lymphocytes ^1, 2^. Although this model was controversial for many years, it is now accepted that mouse embryonic, fetal, and adult hematopoietic stem cells (HSCs) produce distinct populations of immune cells. Indeed, work over the last decade has shown that microglial cells and tissue macrophages originate from yolk sac primitive HSCs ^3, 4^. Similarly, it is established that like B1 lymphocytes ^5^, group 3 innate lymphoid cells (ILC3) and V*γ*5^+^/*γ*6^+^ skin- or genital/urinary tract-resident T cells develop exclusively during embryonic life ^6^. In mice, lymphoid specification is initiated downstream of HSCs in Sca-1^+^c-Kit^+^Flt3^hi^ lymphoid-primed multipotent progenitors (LMPPs) ^7, 8^ which lose myeloid potential as they upregulate the interleukin 7 (IL-7) receptor *α*-chain and differentiate into Lymphoid-primed (LPPs) and then common lymphoid progenitors (CLPs) ^9^. These subsequently segregate into common helper innate lymphoid cell progenitors (CHILPs), thymus settling progenitors (TSPs) and early B cell progenitors (EBPs) ^10^. Despite evidence that mouse T and B cell precursors emerge asynchronously in fetal liver ^11^ and that thymus colonization is ensured by successive waves of T/ILC-restricted and multipotent TSPs ^10^, it is currently unclear whether these changes are exclusively related to different stem/progenitor cell origins or if they could also reflect age-related variations in the lymphoid potential of definitive HSCs ^12^.

The search for ontogeny-related changes in lymphocyte production patterns in humans has long been hampered by limited characterization of lymphoid developmental trajectories and controversies regarding the phenotype and function of lymphoid progenitors ^13^. Since their first identification in postnatal bone marrow (BM) ^14^, extensive study of CD45RA^+^CD10^+^ multi-lymphoid progenitors (referred to as the CD10^+^ MLPs thereafter) has revealed that, despite an intrinsic B-lineage bias ^15^, this population displays multi-lymphoid potential ^16, 17^. However, since CD10^+^ MLPs are only detected after birth ^18, 19^, most current evidence suggests that fetal lymphopoiesis is supported by different and yet uncharacterized MLP subsets. Notably, another subset of CD45RA^+^CD7^+^ MLPs with strong T potential has been isolated from neonatal cord blood ^20, 21^ but their developmental status and biological function remain poorly defined.

We have previously shown that human lymphopoiesis displays a bipartite organization stemming from founder populations of CD127^-^ and CD127^+^ early lymphoid progenitors (ELPs) that differ as to both growth factor dependence and differentiation potential ^18^. Whereas the CD127^-^ ELPs comprise prototypic CD7^+^ T and NK/ILC precursors, their CD127^+^ counterparts are intrinsically biased toward the B lineage. More recently, using an in vivo modeling approach of second trimester fetal hematopoiesis in humanized mice, we have established a map at clonal resolution of human lymphopoiesis (Alhaj Hussen et al., submitted). Our results show that CD127^-^ and CD127^+^ ELPs originate from a novel subset of prototypic CD117^lo^ MLPs lacking both CD7 and CD10 expression and that they are subjected to a differential Flt3L-dependent versus cell-intrinsic regulation. To explore the physiological implications of these findings and reconcile the bipartite model of lymphoid organization ^18^ with earlier studies that categorized lymphoid progenitors on the basis of differential expression of CD7 and CD10 expression ^16^, we subjected lymphoid progenitors from 140 donors between 13 weeks post-conception (PCW) and 87 years postnatal to in-depth immunophenotypic, molecular and functional characterization.

As well as showing that human lymphopoiesis is supported by three waves of embryonic, fetal, and postnatal multi-lymphoid progenitors (MLPs), our data reveal that transition to postnatal life is associated with a previously undiscovered B-lineage shift in the pattern of lymphocyte production.

## RESULTS

### Neonatal CD10^+^ MLPs are intrinsically biased toward CD127^+^ ELPs

To investigate the heterogeneity of lymphoid progenitors circulating in the umbilical cord blood, we developed optimized FACS sorting schemes of CD45RA^+^Lin^-^ hematopoietic progenitor cells (HPCs) by adding ITGB7 and CD127 to a previous six parameter design ^15, 18^. Neonatal CD45RA^+^Lin^-^ HPCs were first partitioned into four ITGB7^-^ or ITGB7^int^ HPC and CD127^-^ or CD127^+^ ELP compartments, each of which was further subdivided according to differential CD7 and CD10 expression (Fig. 1A and Extended Data Fig. 1A-C [step-1 gating procedure]). Hereafter, ITGB7^-^ HPCs will be referred to as lympho-myelo-dendritic progenitors (LMDPs) following a previously established nomenclature ^18^. We then performed B, NK/ILC and T differentiation assays with the resulting 16 cell fractions. As expected, we found hotspots of lymphoid potential within CD127^-^ and CD127^+^ ELPs and confirmed that among these CD7 expression correlates with detection of T/NK-ILC potential (Extended Data Fig. 2). Also consistent with our previous findings ^18^, CD127^+^ ELPs were intrinsically biased toward the B lineage, irrespective of CD7 or CD10 expression.

**Figure 1.**
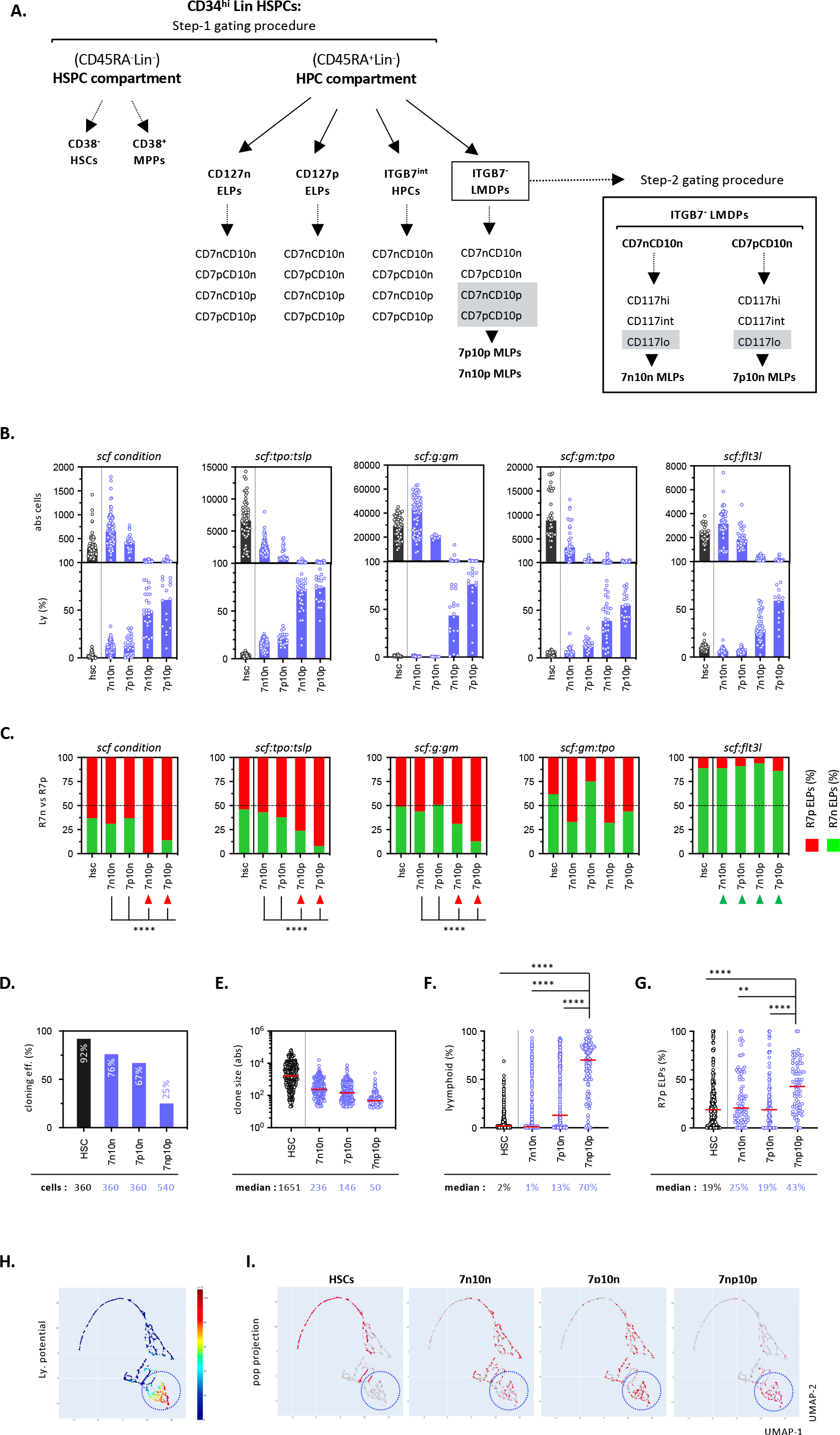
CD10^+^ MLPs are biased towards the CD127^+^ ELPs. **(A).** List and nomenclature of 24 neonatal CD34^+^ HSPCs subsets analyzed in this study. UCB CD34^+^Lin^-^ HSPCs were first subdivided into CD45RA^-^ or CD45RA^+^ compartments. Within CD45RA^-^ compartment, CD38 expression levels distinguished HSCs from MPPs. In Step-1 gating procedure the CD45RA^+^ compartment was first partitioned into ITGB7^-^ LMDPs, ITGB7^int^ HPCs or CD127^-/+^ ELPs (R7np), each of which being further subdivided into CD7 or CD10 negative (n) or positive (p) subsets. In Step-2 gating procedure, CD7^-^CD10^-^ and CD7^+^CD10^-^ LMDPs were further fractionated into CD117^hi/int/lo^ subsets. Lymphoid-restricted CD117^lo^ subsets highlighted in grey are referred to as MLPs in keeping to the current nomenclature. HSC: hematopoietic stem cell, MPP: multipotent progenitor; LMDP: Lympho-Myelo-Dendritic cell Progenitor; HPC: hematopoietic progenitor cell; ELP: Early Lymphoid Progenitor; MLP: multi-lymphoid progenitor. The full gating procedure is presented in Extended Data Fig. 1. **(B, C).** *In vitro* assessment of neonatal LMDP differentiation potentials in bulk diversification assays. **(B)** HSC (grays bars) or LMDPs (blue bars) subdivided based on CD7 and CD10 expression (Step-1 gating procedure) were seeded by 100-cell pools in 96-well plates and cultured for 10 days under the indicated conditions before quantification of *in vitro* differentiated myeloid granulocyte (Gr), monocyte (Mo) and dendritic (DC) precursors, or lymphoid CD127^-/+^ ELPs and CD19^+^ BL (see Extended Data Fig. 3A for the gating procedure); bar plots show absolute cell numbers (upper panels) or percentages of lymphoid cells (lower panels); percentages of lymphoid cells are defined as the sum of the percentages of CD19^+^ BL and CD127^-/+^ ELPs; results are normalized relative to hu- CD45^+^ cells; bars show median values; circles correspond to individual wells. Assessment of statistical significance by the Kruskal-Wallis’ test showed p-values < 0.0001 for all conditions. Quantification of myeloid output is provided as Extended Data Fig. 3B. **(C)** Stacked bar plots show the relative proportions of CD127^-^ (green) and CD127^+^ (red) ELPs. Results are expressed as median percentages from ≥10 individual wells; red and green arrows show polarized ELP production patterns. Assessment of statistical significance was performed with the Mann-Whitney test (CD7npCD10p versus others: p-values < 0.0001 by the Mann-Whitney test [****]). **(D-I).** *In vitro* assessment of neonatal LMDP differentiation potentials in clonal diversification assays. The indicated HSC or LMDP subsets were seeded individually in 96-well plates and cultured for 14 days onto OP9 stroma under the *scf:gm-csf:tpo* condition before quantification of lympho-myeloid output (see Extended Data Fig. 3C for the gating procedure). Positivity threshold for clone detection was se arbitrarily at ≥ 20 cells/clone. Bar and circle plots show **(D**) cloning efficiencies; **(E)** absolute cell numbers per individual clone; **(F)** total lymphoid output or **(G)** relative percentages of CD127^+^ ELPs within each clone normalized relative to total ELPs (positivity threshold for ELP detection was set arbitrarily at ≥ 10 ELPs/clone). For circles plots, red bars show the medians. Corresponding values are indicated below each plot. Note that due to overall similar cloning efficiencies and lymphoid potentials results obtained with CD7^-^CD10^+^ and CD7^+^CD10^+^ MLPs were pooled. Assessment of statistical significance was performed with the Mann-Whitney test (**** p<0.0001; *** p<0.001; ** p<0.01). **(H, I).** Bidimensional projection into a UMAP of lymphoid-containing clones. The Uniform Manifold Approximation and Projection (UMAP) algorithm was applied to the data pooled from all experiments and ancestor cell types (n=2633) with each ancestor cell being projected into the UMAP coordinates. **(H)** Heatmap shows lymphoid potentials defined as percentages of lymphoid cells per clone across all ancestor cells. **(I)** Snapshots show projection of clones derived from the indicated subsets; dashed circles indicate lymphoid-enriched area. Results are pooled from 5 independent experiments.

**Figure 2.**
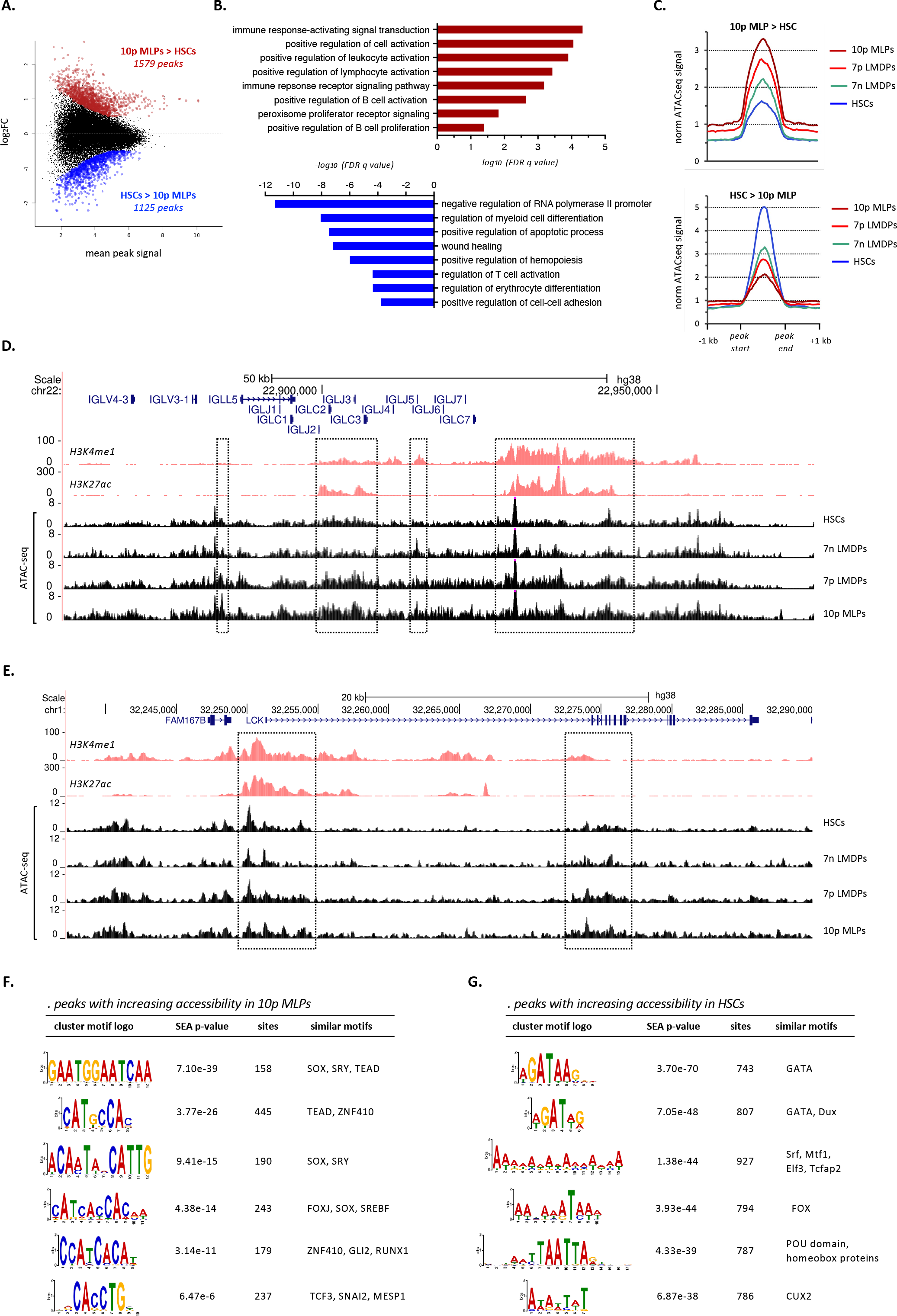
CD10^+^ MLPs are epigenetically primed toward the B lineage. **(A).** Mean-difference (MD) plot of 46531 total ATAC-seq peaks showing log2(fold change) of ATAC-seq peak signal in CD10^+^ (10p) MLPs relative to signal in HSCs against mean signal of each peak; peaks increasing (FDR q<0.05, log2FC>0.5) or decreasing (FDR q<0.05, log2FC<-0.5) in CD10^+^ MLPs relative to HSCs are indicated in red (1579 peaks) and blue (1125 peaks) respectively. **(B).** Selected Gene Ontology (Biological Process) categories enriched (FDR q<0.05) in genes associated with peaks of increased (cayenne) or decreased (blue) accessibility in CD10^+^ MLPs relative to HSCs. The full list of enriched categories is provided in Table S1. **(C).** Cumulative normalized ATAC-seq signal at peaks increasing (1579 peaks; top panel) or decreasing (1125 peaks, bottom panel) in CD10^+^ MLPs relative to HSCs; shown is the mean cumulative signal observed at these peak sets in the indicated populations. **(D, E).** UCSC Genome Browser snapshots showing changes in chromatin accessibility observed in the indicated populations at *cis*-elements within the **(D)** IG lambda and **(E)** LCK loci (from GO category: Immune response-activating signal transduction); gene annotation (Gencode V38) is shown at top followed by Encode tracks demonstrating enhancer-associated histone modifications (H3K4me1 and H3K27ac) in the GM12878 lymphoblastoid cell line. Regions whose accessibility increases during lymphoid specification are highlighted (dashed boxes). **(F, G).** Results of motif enrichment analysis (XSTREME) showing the 6 most enriched motif clusters detected in peaks of increased accessibility in **(F)** 10p MLPs or **(G)** HSCs; shown for each cluster is the significance of enrichment (Simple Enrichment Analysis E-value), the number of times that the motif is detected in the test peak set (sites), and similar known motifs detected in the TomTom dataset. Abbreviations: 10p: CD10^+^CD7^-^; 7p: CD10^-^CD7^+^, 7n: CD10^-^CD7^-^.

Since only marginal lymphoid potential was detected within the more immature LMDP and ITGB7^int^ compartments, we next investigated whether this was due to a lack of lymphoid competence or to the limited sensitivity of standard differentiation assays. To address this, LMDPs fractionated as above were seeded in multi-lineage diversification assays allowing simultaneous generation of granulomonocytic, dendritic and lymphoid precursors (Extended Data Fig. 3A). Cultures were supplemented with *scf* only or with cytokine cocktails promoting lymphoid (*scf:tpo:tslp*), multi- lineage (*scf:flt3l* or *scf:gm:tpo*) or myeloid (*scf:g:gm*) differentiation (Alhaj Hussen et al., submitted). Consistent with an earlier report ^16^, the CD10^+^ fractions expanded poorly and generated a majority of lymphoid cells (Fig 1B and Extended Data Fig. 3B). Except for the *scf:flt3l* condition that induced a complete lymphoid shift toward CD127^-^ ELPs, the CD10^+^ fractions were markedly biased towards the CD127^+^ ELPs, with no consistent difference being noted with respect to CD7 co-expression (Fig. 1C). This contrasts with their CD10^-^ counterparts that expanded robustly, showed mixed lympho-myeloid potentials, and displayed balanced CD127^-^ and CD127^+^ ELP output. Single-cell differentiation assays (Extended Data Fig. 3C) confirmed that the CD10^+^ fractions have reduced cloning efficiency, undergo limited expansion and that they mostly generate lymphoid cells with a high proportion of CD127^+^ ELPs (Fig. 1D-I and Extended Data Fig. 3D-F). In contrast, their CD10^-^ counterparts proved highly heterogeneous including multipotent as well as already myeloid or lymphoid lineage-specified progenitors with, here again, balanced lymphoid output.

**Figure 3.**
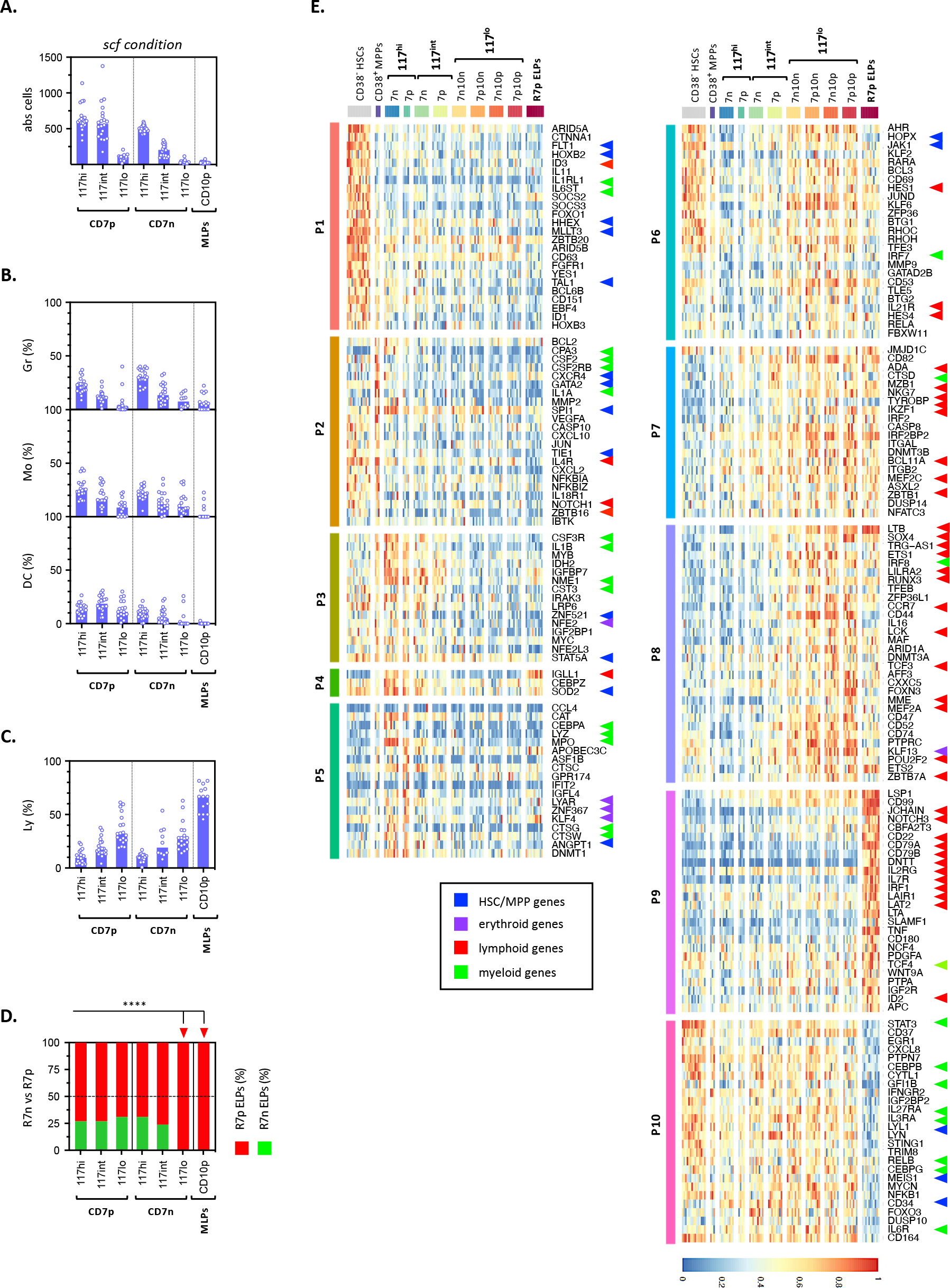
Molecular and functional characterization of neonatal MLPs. **(A-D).** Functional characterization of CD117^hi/int/lo^ LMDP subfractions in bulk diversification assays. CD7^-^ or CD7^+^ LMDPs were partitioned into CD117^hi/int/lo^ fractions (Extended Data Fig. 1C: step-2 gating procedure) and seeded as above under the *scf* condition before quantification of cell growth and lineage output. Bar plots show **(A)** absolute cell numbers; **(B)** percentages of myeloid granulocyte (upper panel), monocyte (medium panel) or dendritic (lower panel) precursors; **(C)** percentages of lymphoid cells. Bars indicate medians; circles correspond to individual wells; assessment of statistical significance was performed with the Kruskal-Wallis test (****p <0.0001 for all conditions). Stacked bar plot shows **(D)** the relative proportions of CD127^-^ (green) and CD127^+^ (red) ELPs; results are expressed as median percentages from ≥10 individual wells; red arrows show statistically significant differences (CD117^lo^CD7^-^10^-^ or CD10^+^ versus others; **** p < 0.0001 by the Mann-Whitney test). **(E).** The indicated neonatal subsets were analyzed mini-RNAseq. Heatmaps showing expression of 209 selected DEGs across 10 clusters (P1-10). Gene expression values are log-transformed, normalized and scaled; colored arrows show lineage- or subset-specific genes. See also Table S2 for the distribution of the 1,773 DEGs within clusters P1-10.

These data confirm that expression of CD10 by neonatal LMDPs is indicative of lymphoid specification and an intrinsic differentiation bias toward the generation of CD127^+^ ELPs. In keeping with the current nomenclature ^16^, CD10^+^ LMDPs are hereafter referred to as CD10^+^ MLPs.

### Neonatal CD10^+^ MLPs are epigenetically primed toward the B lineage

To investigate the molecular events underlying intrinsic B-lineage bias of neonatal CD10^+^ MLPs, we carried out ATAC-seq experiments to compare their chromatin accessibility profile with that of HSCs, and of CD7^-^CD10^-^ and CD7^+^CD10^-^ LMDPs (Extended Data Fig. 4A). Consistent with recent findings that HSCs exhibit a generally open chromatin structure ^22^, the highest number of peaks was observed in HSCs (40763 peaks, FDR<0.05). Peak number decreased progressively in CD7^-^CD10^-^ (35457 peaks) and CD7^+^CD10^-^ LMDPs (26910 peaks), and the lowest number of peaks was detected in CD10^+^ MLPs (19291 peaks) consistent with a gradual extinction of alternative lineage potentials during lymphoid specification (Extended Data Fig. 4B).

**Figure 4.**
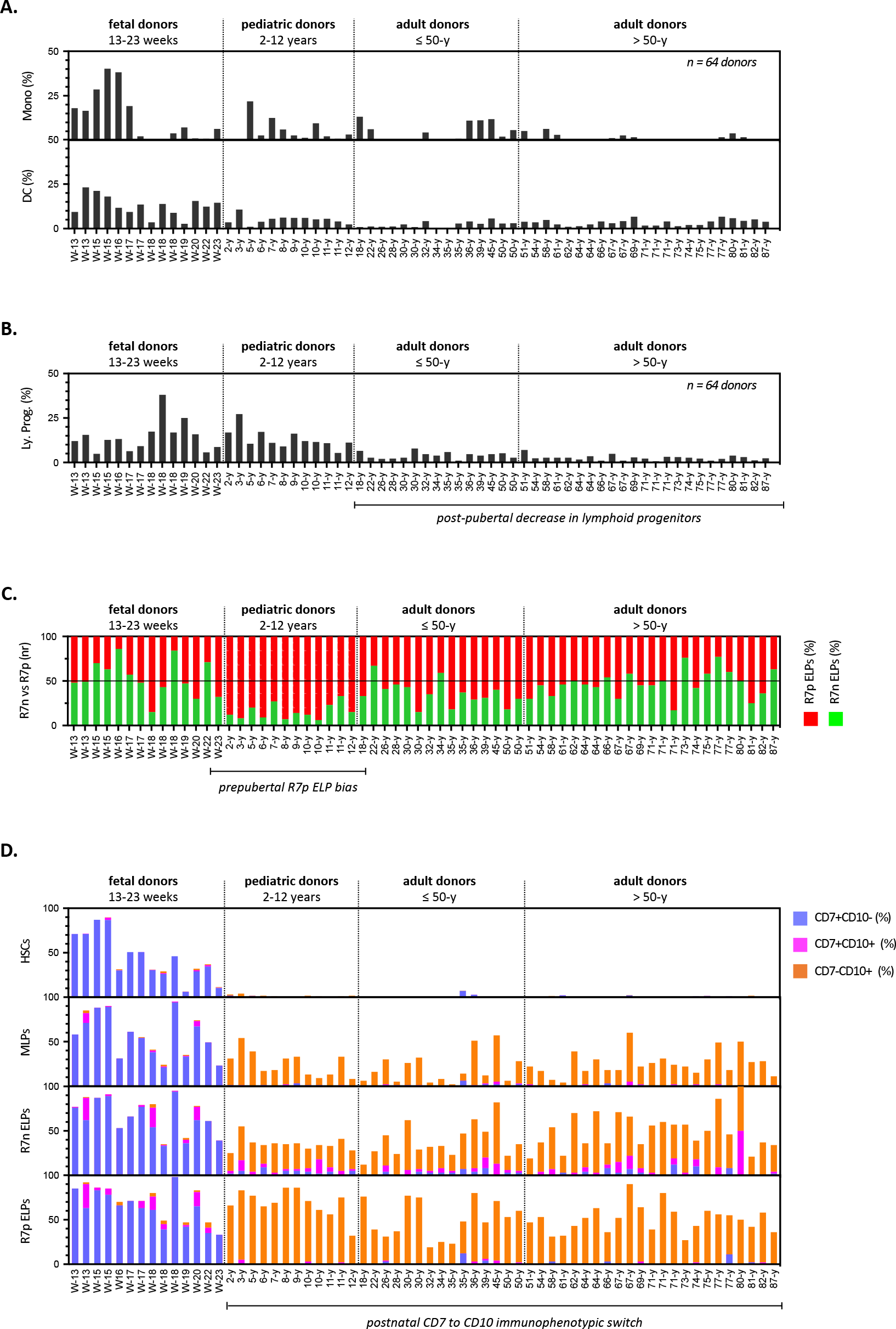
Transition to postnatal life coincides with a B-lineage shift of lymphopoiesis. **(A, B).** Dynamics of myeloid **(A)** monocyte (upper panel) and dendritic (lower panel) precursors, or of **(B)** lymphoid progenitors in the BM of 14 fetal and 50 postnatal donors. Gates were set as described in Extended Data Fig. 7 (Gating Strategy-1); results are normalized according to total CD34^hi^Lin^-^ HSPCs. **(C).** Stacked bar plots show the relative proportions of CD127^-^ (green) and CD127^+^ (red) ELPs in the same donors; results are normalized relative to total ELPs. **(D).** Stacked bar plots show age-dependent variations in CD7 or CD10 expression across HSC, MLP, CD127^-^ or CD127^+^ ELP compartments in the same donors; results are expressed as relative percentages within each subset.

Subsequent analysis was carried out on a master set of 46531 peaks, comprised of all peaks present in at least one population. To establish the profile of chromatin changes associated with lymphoid commitment, we initially compared the accessibility at these peaks between HSCs and CD10^+^ MLPs and identified 1579 peaks with increased accessibility and 1125 peaks with decreased accessibility (FDRq<0.05, log2FC>0.5) during lymphoid commitment (Fig. 2A). Relative to the complete peak set, peaks which change during lymphoid commitment are enriched at intronic and gene-distal sites (Extended Data Fig. 4C), consistent with previous findings that hematopoietic lineage commitment is primarily dictated by gene-distal enhancer elements ^23^. Genes associated with peaks of increasing accessibility in CD10^+^ MLPs were enriched for pathways of immune receptor signaling, B cell activation and expansion, confirming their lymphoid-committed status and B lineage bias (Fig. 2B and Table S1). In contrast, genes associated with sites of decreasing accessibility were enriched for general cellular processes (transcription, apoptosis, cell adhesion) as well as the regulation of myeloid, T lymphoid and erythroid pathways. Pileup analysis of accumulated ATAC signal confirmed that these changes are partially detected in CD7^-^CD10^-^ LMDPs and increase further in CD7^+^CD10^-^ LMDPs (Fig. 2C and Extended Data Fig. 4D, E). Representative examples of gradually acquired accessibility at regulatory elements associated with lymphoid genes (IGL, LCK, TNFAIP3) and their gradual decline at genes important for alternative cell fates (for example the erythroid-determining factor NFE2) are shown (Fig. 2D, E and Extended Data Fig. 4F, G). Consistent with these results, analysis of DNA motifs in regions of increased accessibility in CD10^+^ MLPs revealed strong enrichment for motifs recognized by the B-lymphoid transcription factors SOX4, TEAD1/TEF1 or TCF3 ^24^, while peaks of decreasing accessibility were most enriched for motifs recognized by the GATA family of transcription factors (Fig. 2F, G).

These results confirm that CD10^+^ MLPs are epigenetically primed toward the B lineage and demonstrate that the molecular changes associated with lymphoid specification are progressively acquired.

### Functionally distinct CD7^-/+^CD10^-/+^ MLP subsets circulate in neonatal blood

We have recently shown that the lymphoid potential of fetal LMDPs can be defined based on decreasing CD117 expression (Alhaj Hussen et al., submitted). To determine whether this also applies to neonatal LMDPs, cord blood CD7^-/+^CD10^-/+^ LMDP subsets were stratified according to CD117 levels (Extended Data Fig. 1C [step-2 gating procedure]). As expected, we found that CD10^+^ MLPs express uniformly low levels of CD117. Since CD7^+^ or CD7^-^ LMDPs express variable levels of CD117, they were further fractioned on this basis and seeded as above in diversification assays. Culture under *scf* or *scf:tpo:tslp* conditions confirmed that downmodulation of CD117 coincides here again with reduced proliferation, decreasing myeloid potential and a corresponding increase in lymphoid output (Fig. 3A-D, Extended Data Fig. 5A-D). Furthermore, analysis of ELP production patterns revealed that, conversely to their CD7^-^ or CD10^+^ counterparts that are markedly biased toward CD127^+^ ELPs, the CD117^lo^CD7^+^ LMDPs give rise to both CD127^-^ and CD127^+^ ELP output. Single-cell cultures confirmed that CD117^lo^CD7^+^ LMDPs display limited cloning efficiency, reduced expansion and mostly generate lymphoid cells among which CD127^-^ ELPs predominate (Extended Data Fig. 5E-H).

**Figure 5.**
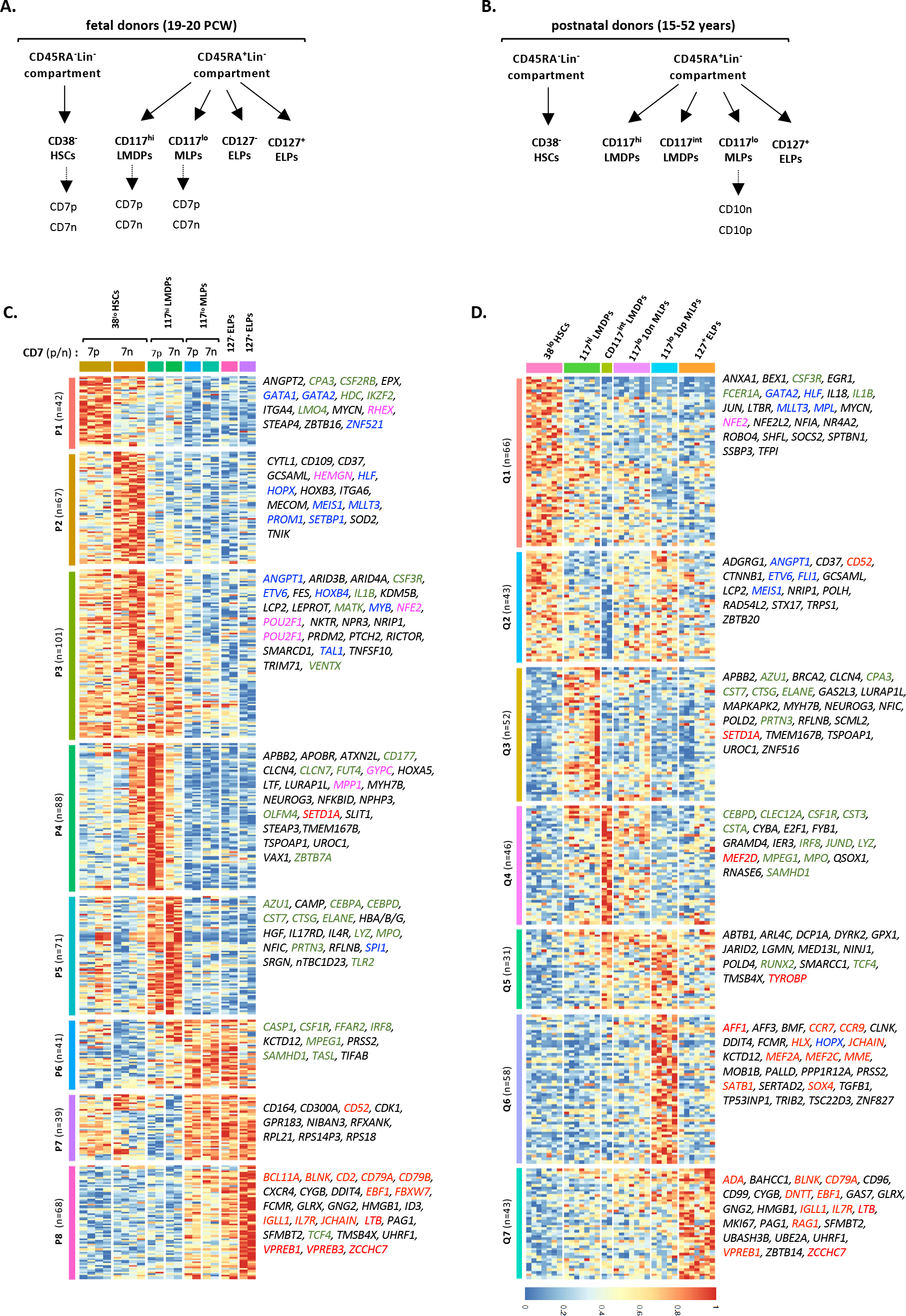
Transcriptional profiling of lymphoid progenitors from fetal or adult BM. The indicated HSPC subsets were sorted from **(A)** 2 fetal (19-20 PCW) or **(B)** 4 postnatal donors (15- 52 years) as described in Extended Data Fig. 7 (Gating Strategy-1) and processed for transcriptional profiling by mini-RNAseq. Heatmaps show expression of **(C)** 517 DEGs between 8 fetal subsets across 8 clusters (P1-8), and of **(D)** 339 DEGs between 6 postnatal subsets across 7 clusters (Q1-7). Gene expression values are log-transformed, normalized, and scaled; right panels correspond to selected short lists of biologically significant cluster-specific genes. Functionally annotated HSC/MPP (blue), erythroid (purple), myeloid (green) or lymphoid (red) genes are indicated.

Collectively, these data indicate that distinct subsets of CD7^+^CD10^-^, CD7^-^CD10^-^, and CD7^-/+^CD10^+^ MLPs with balanced or B lineage-biased lymphoid differentiation potentials circulate in the neonatal cord blood.

### Neonatal CD7^-/+^CD10^-/+^ MLPs display overlapping gene signatures

To characterize the changes in gene expression associated with lymphoid specification and search for transcriptional correlates of the differences in the pattern of ELP production observed between CD7^-/+^CD10^-/+^ MLPs, 11 neonatal HSPC populations were subjected to transcriptional profiling using ultra-low-input mini-RNA-sequencing (Extended Data Fig. 5I).

Principal component analysis (PCA) showed co-clustering of all MLP subsets and confirmed that downstream CD127^+^ ELPs are more distantly related (Extended Data Fig. 5J). Within CD117^int/hi^ LMDPs, PCA also allowed distinction between the CD7^-^ and CD7^+^ cell fractions, with the latter overexpressing Notch target genes (*HES1*, *FBXW5*, *CDCA7*) (data not shown). A total of 55 pairwise comparisons between the indicated subsets identified 1,773 differentially expressed genes (DEG) distributed in 10 clusters (P1-10) (Extended Data Fig. 5K and Tables S2A, B). Functional analysis of gene enrichment patterns and cluster distribution recapitulated the hematopoietic hierarchy with the most immature CD38^-^ or CD38^+^CD45RA^-^ subsets overexpressing genes involved in stem cell maintenance (P1: *FLT1*, *HOXB2*, *TAL1*, *HHEX*) and/or early hematopoietic differentiation (P2: *GATA2*, *SPI1*, *TIE1*, *CSF2RB*, *CXCR4*) (Fig. 3E). Consistent with granulomonocytic specification, CD117^hi^ LMDPs upregulated *CSF3R*, *CEBPA*, *LYZ*, *MPO*, *CTSG* and *CTSW* transcripts (P3-5). As expected, onset of lymphoid priming was detected within the CD117^int^ LMDP fractions (P6-7) and coincided with overexpression of Notch target genes *HES1* or *HES4*, acquisition of *IL21R* and upregulation of transcription factors (TFs) controlling early lymphoid specification (I*KZF1*, *BCL11A*, *ZBTB1*) or dendritic cell differentiation (*IRF8*). A second wave of lymphoid TFs (*ETS1*, *RUNX3*, *TCF3*, *SOX4*, *ZBTB7A*) was detected in downstream CD117^lo^ MLPs (P8). Gene expression in CD117^lo^ MLPs was similar irrespective of CD7 or CD10 expression. Finally, consistent with a more advanced differentiation status, the highest expression levels of *IRF1*, *ID2*, *TCF4* and *Notch3*, as well as of BCR components (*CD22*, *CD79A*, *CD79B*, *LAT2*) were detected in CD127^+^ ELPs (P9), that no longer expressed stem/progenitor cell-associated TFs (*LYL1*, *MEIS1*) or myeloid lineage (*STAT3*, *CEBPB*, *CEBPG*, *GFI1B*, *IL3RA*, *IL6R*) genes (P10).

Regulatory network analysis including transcription factors (TFs) upregulated in neonatal MLPs (P6-8) and their potential target genes confirmed the role played by MEF2-family transcription factors during lymphoid specification ^25^. This also revealed three additional densely connected nodes represented by the HOX-family tumor suppressor CUX1 ^26^, as well as by the TFEB activator of lysosomal activity ^27^ and the FOXN3 cell-cycle regulator ^28^, which suggests that following egress of into the blood these MLPs upregulate a quiescence program (Extended Data Fig. 6A).

**Figure 6.**
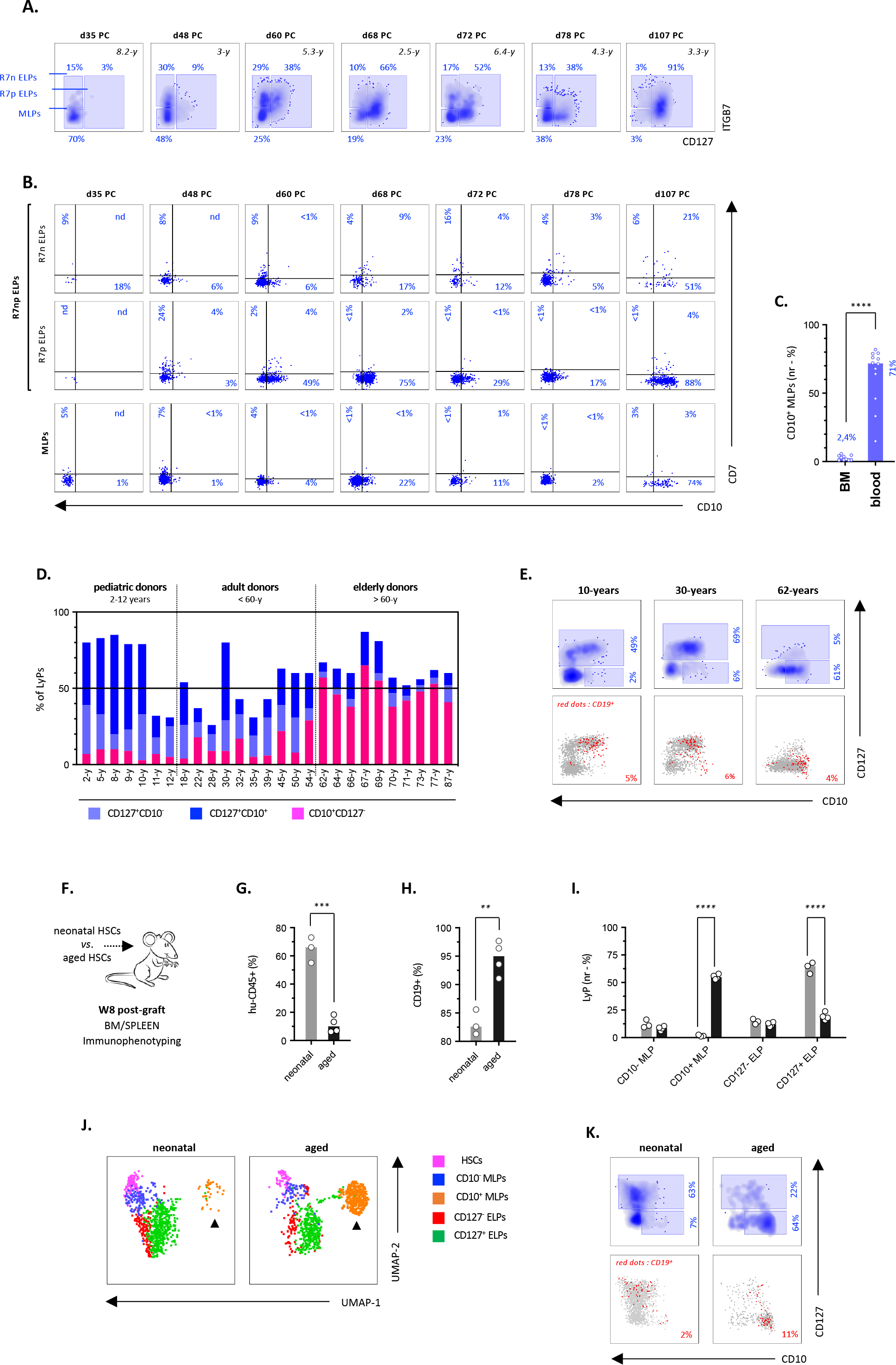
Kinetics of CD10^+^ MLPs and CD127^+^ ELPs in postnatal donors. **(A, B).** Immunophenotypic profiling of lymphoid progenitors in the BM of pediatric patients (2.5 to 8.2 years) between 35 to 107 days after treatment for AML or ALL. Gates were set as described in Extended Data Fig. 7 (strategy 1). **(A)** Upper density plots show the subdivision of lymphoid progenitors into MLP, CD127^-^ or CD127^+^ ELP subsets; **(B)** lower dot plots show the expression of CD7 and CD10 by the corresponding subsets. Percentages are indicated. **(C).** Quantification of blood and BM CD10^+^ MLPs in 22 pediatric donors (blood, n=12; BM, n=10) between 2 and 12-years. Results are normalized relative to total CD34^+^CD45RA^+^Lin^-^ HPCs. Bars indicate medians; circles correspond to individual donors; assessment of statistical significance was performed with by the Mann-Whitney test) (****p <0.0001 for all conditions). **(D).** Dynamics of BM CD10^+^ MLPs and CD127^+^ ELPs in 27 postnatal donors between 2 and 87 years; gates were set as described in Extended Data Fig. 7 (Gating Strategy-3). Stacked bar plot show percentages of the indicated subsets. **(E).** Immunophenotypic mapping of B cell differentiation pathways in 3 representative BM donors. Upper density plots showing age-related changes in BM CD10^+^ MLPs and CD127^+^ ELPs; lower dot plots overlays show projection of CD19^+^ cells within the corresponding subsets; percentages are indicated. **(F-K).** Lymphoid reconstitution in mice xenografted with HSCs from elderly donors. **(F)** Experimental design: NSG mice reconstituted with 2,5 x 10^5^ HSCs from two donors aged 65 and 80-years (2 mice/donor) or from the UCB (n=3) were sacrificed 8 weeks after grafting. Immunophenotypic profiling of hu-CD45^+^ BM and spleen cells was performed by spectral cytometry using a 35-antibody panel. Bar plots show percentages of **(G)** hu-CD45^+^ cells (normalized relative to total BM mononuclear cells); **(H)** CD19^+^ B lymphocytes (normalized relative to hu-CD45^+^ cells) or of **(I)** the indicated subsets (normalized relative to total Lymphoid progenitors) (see also Extended Data Fig. 7 Strategy-1 for the gating procedure); circles correspond to individual mice; bars indicate medians; assessment of statistical significance is based on the unpaired t-test (**; p<0.01; ***; p<0.001). (**J)** Immunophenotypic profiling of hu-CD34^hi^ HSPCs in mice xenografted with HSCs from neonatal (left panel) or aged (right panel) donors; gates were set as described in Extended Data Fig. 7 (Strategy- 1); UMAP shows projection of the indicated subsets; FACS data are pooled from individual mice. **(K)** Immunophenotypic mapping of B-cell differentiation pathways in the same xenografted mice; upper density plots showing ontogeny related changes in CD10^+^ MLPs and CD127^+^ ELPs; lower dot plot overlays show projection of CD19^+^ cells within the corresponding subsets; percentages are indicated.

In conclusion, these results show that, irrespective of CD7 or CD10 expression, neonatal MLPs display overlapping lymphoid transcriptional signatures, consistent with the idea that the differences in ELP production pattern reported above are conditioned at epigenetic level.

### Differential CD7 and CD10 expression identifies embryonic, fetal, and postnatal MLP subsets

Inasmuch as earlier studies provided evidence that CD7 is expressed preferentially during the fetal period ^20, 29^, we next hypothesized that rare CD7^+^ MLPs circulating in the neonatal blood could have an embryonic or fetal origin. To address this question, BM CD34^+^ HSPCs from 14 fetal (13-23 PCW) and 50 postnatal (2-87 years) donors were subjected to optimized immunophenotypic profiling (Extended Data Fig. 7 [Strategy-1]). As well as showing a predominance of monocyte and dendritic precursors in the BM of fetal donors before 16 PCW, these analyses found that lymphoid progenitors reach maximum levels by 18-20 PCW and remain at stable levels until puberty, after which they gradually decline to represent ≤ 3% of total HSPCs after 60 years (Fig. 4A, B). Further dissection of lymphoid architecture revealed that whereas CD127^-^ and CD127^+^ ELPs are produced in comparable amounts throughout the fetal period, a clear bias toward CD127^+^ ELPs is observed immediately after birth and persists until puberty (Fig. 4C). Subsequent assessment of CD7 and CD10 expression patterns showed that whereas early in development FL and BM CD34^+^ HSPCs homogeneously express CD7 irrespective of differentiation stage, beyond first trimester gestation the percentages of CD7^+^ HSPCs gradually decline until birth (Fig. 4D and Extended Data Fig. 8A). Despite restriction to the lymphoid lineage, the pattern of expression of CD10 also showed ontogeny-related variations. In contrast to the fetal period during where it remains confined to CD19^+^ B-cell precursors ^18, 19^, after birth the expression of CD10 is acquired at earlier differentiation stages starting at the stage of MLPs. Further immunophenotypic profiling of CD34^+^ HSPCs in the blood of 47 fetal or neonatal donors between 19 PCW and 77-day postnatal confirmed that the immunophenotypic switch from CD7 to CD10 on lymphoid progenitors takes place within the first few days after birth (Extended Data Fig. 8B).

**Figure 7.**
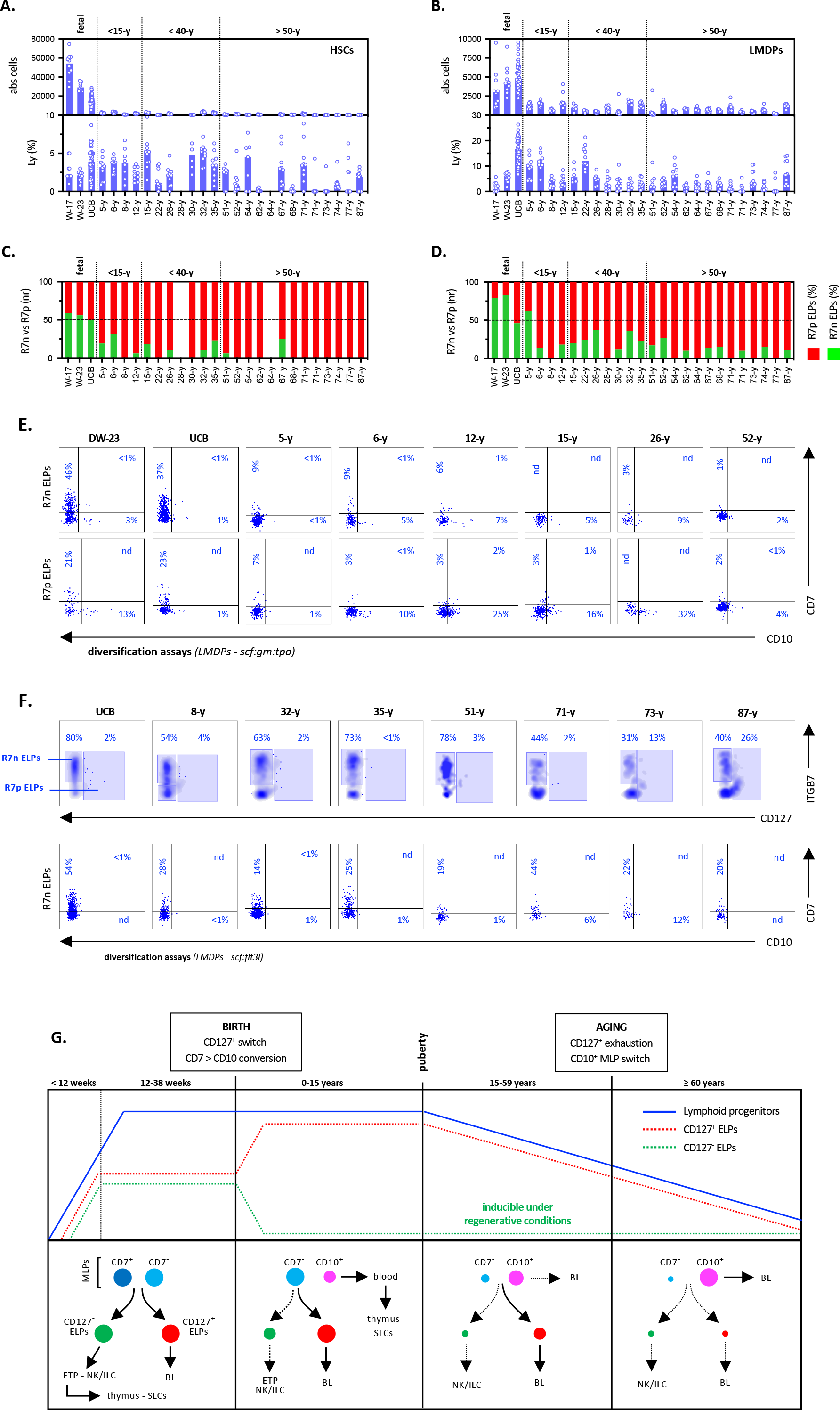
Ontogeny-related changes in lymphoid potential are orchestrated at the level of HSCs. **(A-D).** In vitro assessment of the lymphoid potential of fetal and postnatal CD34^+^ HSPCs. The indicated HSC (left panels) or LMDP (right panels) fraction isolated from the UCB or BM of 26 donors between development week-17 and 87 years (Extended Data Fig. 7: Gating Strategy-3) were cultured for 10 days under the *scf:tslp:tpo* condition before quantification of lymphoid outputs. Bar plots show absolute cell numbers (upper panels) or percentages of lymphoid cells (lower panels) in **(A)** HSC or **(B)** LMDP cultures; bars indicate medians; circles correspond to individual wells. Stacked bar plots show the relative proportions of CD127^-^ (green bars) and CD127^+^ (red bars) ELPs in **(C)** HSC or **(D)** LMDP cultures; bars correspond to median percentages of ≥10 individual wells. **(E).** Dot plots showing age-dependent variations in CD7 and CD10 expression by *in vitro* differentiated CD127^-^ or CD127^+^ ELPs. Cells were sorted, cultured, and analyzed as above; data are from 10 concatenated wells. **(F).** Effect of Flt3L on CD127^-^ or CD127^+^ ELP differentiation (upper panel) and CD7 or CD10 expression (lower panels). LMDPs sorted from the UCB or the BM of 8 postnatal donors were cultured under the *scf:flt3l* condition and analyzed as above; upper density plots show CD127^-^ or CD127^+^ ELPs; lower dot plots show CD7 and CD10 expression; percentages are indicated. **(G).** Model recapitulating the developmental transitions in lymphopoiesis occurring across ontogeny. Upper panels show the dynamics of lymphoid progenitors (plain blue line), as well as of CD127^-^ (dashed green line) or CD127^+^ ELPs (dashed red line). Lower panels show age-related changes in lymphoid differentiation pathways: plain and dashed arrows indicate predominant and accessory differentiation pathways; circle diameters correspond to the estimated size of the corresponding cell subsets. Dark blue circles: embryonic CD7^+^ MLPs; light blue circles: fetal CD7^-^ MLPs; purple circles: postnatal CD10^+^ MLPs); green circles: CD127^-^ ELPs; red circles: CD127^+^ ELPs. SLCs: secondary lymphoid organs.

Collectively, these results show that the transition to postnatal life coincides with a global B-lineage shift in lymphopoiesis characterized by a prepubertal increase in production of CD127^+^ ELPs accompanied by an immunophenotypic CD7 to CD10 conversion of lymphoid progenitors which takes place around birth and is maintained throughout postnatal life. Further, our data provide evidence that CD7^+^CD10^-^, CD7^-^CD10^-^, and CD7^-^CD10^+^ MLPs correspond to embryonic, fetal, and postnatal MLPs, respectively.

### Fetal lymphoid progenitors retain expression of myeloid lineage genes

To search for ontogeny-related changes in lymphoid transcriptional signatures, mini-RNA-sequencing was performed on 14 HSPC subsets from 2 fetal (19-20 PCW) and 4 postnatal (15-52 years) BM donors (Fig. 5A, B and Extended Data Figure 7 [Strategy-1]). Consistent with a previous report ^30^, global comparison between fetal and postnatal CD34^+^ HSPC subsets (1,034 DEGs) showed *LIN28B* overexpression and enrichment of cell-cycle, protein synthesis and oxidative phosphorylation pathways in the fetal subsets versus stress- or inflammatory-response pathways in their postnatal counterparts (Extended Data Fig. 9A, B and Tables S3A, B).

Transcriptional profiling of fetal HSPCs identified 517 DEGs distributed in 8 clusters (P1-8) (Fig. 5C & Extended Data Fig. 9; Table S4A, B). Consistent with a more primitive developmental status, gene enrichment analyses revealed that the CD7^+^ HSCs (CD45RA^-^CD38^-^) are enriched with genes involved in stem cell activation/proliferation (*MYCN*, *GATA1*, *GATA2*, *ZBTB16*) and eosinophil/basophil or mast cell differentiation (*LMO4*, *IKZF2*, *CPA3*, *HDC*) (P1), while their CD7^-^ counterparts upregulate markers of postnatal HSCs (*ITGA6/CD49F*, *Prom1/CD133*) and regulators of stem cell self-renewal (*HLF*, *HOPX*, *MEIS1*, *MLLT3*, *HOXB3, SOD2*) (P2). Despite these differences, CD7^+^ and CD7^-^ HSCs shared expression of genes involved in multilineage priming (*VENTX*, *MECOM*, *ARID3B, ARID4A, POU2F1*) and erythroid (*TAL1*, *NFE2*) or myeloid (*CSF3R*, *IL1B*) differentiation (P3). Downstream CD7^+^CD117^hi^ HPCs unexpectedly showed enrichment in genes involved in lung, kidney, cardiovascular or brain development (*HOXA5*, *NPHP3*, *VAX1*) (P4), and shared with their CD7^-^ counterparts expression of granulocyte genes (P5: *CEBPA*, *MPO*, *ELANE*). These analyses also showed that, in contrast to their neonatal counterparts, fetal MLPs display a composite gene signature characterized by combined enrichment in monocyte/dendritic (*CSF1R*, *IRF8*, *MPEG1*, *SAMHD1*) and lymphoid (*LTB*, *CD2*, *ID3*, *IL7R*, *BLNK*, *VPREB1/3*) lineage genes, the latter’s being further upregulated in downstream CD127^-^ or CD127^+^ ELPs (P6-8). Again, no difference in gene expression was noted between CD7^-^ or CD7^+^ MLP subfractions.

Gene enrichment and functional analyses of adult CD34^+^ HSPCs (339 DEGs; Fig. 5D and Extended Data Fig. 9D; Table S5A, B) confirmed overexpression of genes associated with stress inflammatory response (*JUN*, *NFE2L2*, *IL1B*, *EGR1*) and resistance to apoptosis (*AVP*, *CLU*, HSBP1, *PRDX1*) in the HSC compartment (Q1-2) and found the expected granulocyte (Q3) or monocyte/dendritic (Q4-5) gene signatures within CD117^hi^ and CD117^int^ LMDPs. In contrast to their fetal counterparts, adult CD117^lo^CD10^-^ MLPs downregulated most myeloid genes while showing limited evidence of lymphoid priming that became obvious only in CD10^+^ MLPs. Here again, CD10^+^ MLPs and CD127^+^ ELPs displayed the expected multi-lymphoid (*TYROBP*, *MEF2A/C*, *SATB1*, *SOX4*, *CCR7/9*) (Q6) or B lineage-biased gene signatures (Q7). Subsequent regulatory network analysis of the TFs and their potential target genes overexpressed by fetal and postnatal MLPs (Extended Data Fig. 6B) disclosed 2 densely connected nodes corresponding to SATB1 ^31^ and MEF2-family transcription factors ^25^, an indication that their role in lymphoid specification is conserved across ontogeny.

Collectively, these data show that, conversely to their postnatal counterparts, proliferating fetal lymphoid progenitors retain expression of monocyte or dendritic lineage genes. Further, they indicate that in the adult BM lymphoid commitment coincides with CD10 upregulation.

### CD10^+^ MLPs circulate in high numbers in the blood of prepubertal donors

To further investigate the developmental status and biological function of postnatal CD10^+^ MLPs, we next monitored lymphoid regeneration in 7 pediatric patients (2 to 8-years) between 35 and 107 days after treatment by chemotherapy for acute lymphoblastic or myeloblastic leukemia (Extended Data Fig. 7 [strategy-1]). Immunophenotypic profiling of lymphoid progenitors in these patients confirmed that CD127^-^ and CD127^+^ ELPs emerge sequentially within the first two months after treatment (Alhaj Hussen et al., submitted), with the latter appearing to accumulate throughout the regenerative period (Fig. 6A). These analyses also revealed that only a minority of newly produced ELPs express CD7 which confirms that, even under regenerative conditions, postnatal HSCs have a decreased capacity to enter the NK/ILC/T differentiation pathway (Fig. 6B upper panels). We also found that from the second month after chemotherapy, a large proportion of newly produced CD127^+^ ELPs upregulate CD10, indicating their progression along the B-cell differentiation pathway. However, to our surprise, within the first three months of chemotherapy, only rare CD10^+^ MLPs were detected in the BM of these patients. Normal levels of CD10^+^ MLPs were only observed around day 100 post-chemotherapy (Fig. 6B lower panel). As well as indicating that under regenerative conditions CD10^+^ MLPs are late products of lymphoid diversification, these data show that they are not obligatory precursors of CD127^-^ and CD127^+^ ELPs after birth. Since CD10^+^ MLPs have been previously proposed as postnatal thymus colonizers ^32, 33^, we next compared their distribution between blood and BM compartments in 22 pediatric donors between 2-12 years (Fig. 6C). These analyses revealed that percentages of CD10^+^ MLPs are 30-fold higher in blood than BM indicating that during the prepubertal period these MLPs display a selective capacity to enter the circulation possibly to reach the thymus or secondary lymphoid organs.

Altogether, these results indicate that immediately after birth and throughout the prepubertal period lymphoid architecture is characterized by a functional tripartition between CD127^-^ and CD127^+^ ELPs and circulating CD10^+^ MLPs. They also suggest that before puberty CD10^+^ MLPs contribute only marginally to BM lymphopoiesis.

### Identification of a direct CD10^+^ MLP based B-cell differentiation pathway in the elderly

To further investigate the developmental relationships between CD10^+^ MLPs and CD127^+^ ELPs, lymphoid progenitors from 27 donors between 2 and 87 years were subjected to optimized immunophenotypic fractionation (Extended Data Fig. 7 [Strategy-2]). This showed that whereas CD127^+^ ELPs largely predominate during the first five decades of life, CD10^+^ MLPs become the major lymphoid-progenitor subset after 60 years of age, thus arguing for a late shift in B-cell differentiation pathways (Fig. 6D). Subsequent mapping of B-cell differentiation trajectories confirmed that in younger donors CD127^+^ ELPs emerge from the CD10^-^ MLPs and sequentially upregulate CD10 and CD19 (Fig. 6E). This contrasted with elderly donors in which B-cell differentiation stems directly from CD10^+^ MLPs, largely bypassing the CD127^+^ ELPs.

To examine whether this late transition in the B-lineage differentiation paths is developmentally imprinted, NOD-*Scid-IL2rg*^null^ (NSG) mice were transplanted with CD45RA^-^CD38^lo^ HSCs isolated from the BM of 65- and 80-year-old healthy donors and sacrificed two months later (Fig. 6F). Consistent with decreased proliferation of postnatal HSCs, hu-CD45^+^ cells represented ≤10% of BM cells in mice transplanted with aged HSCs, compared to ≥60% for those reconstituted with neonatal HSCs (Fig. 6G). Also, as expected, higher percentages of BM CD19^+^ B lymphocytes were also detected in mice transplanted with aged HSCs (Fig. 6H). Spectral immunophenotyping of CD34^hi^ HSPCs confirmed that whereas only rare CD10^+^ MLPs are detected in mice transplanted with neonatal HSCs, these account for the majority of lymphoid progenitors in those mice reconstituted with aged HSCs (Fig. 6I, J and Extended Data Fig. 10A upper panels). Again, the percentages of CD10^+^ MLPs and CD127^+^ ELPs varied in inverse proportions, reinforcing the idea that with age these populations are subjected to a differential regulation. Further immunophenotyping confirmed that in mice transplanted with aged HSCs B-cell differentiation undergoes a clear switch toward the CD10^+^ MLP pathway (Fig. 6K). Interestingly, detection of few CD7^+^ ELPs in the BM of mice reconstituted with HSCs from older donors is concordant with clinical data showing de novo production of NK cells and ILCs in patients transplanted with adult HSCs ^34, 35^ (Extended Data Fig. 10A lower panels). Finally, consistent with the established age-related myeloid bias in HSC differentiation potential ^36^, analysis of mature cellular populations found that the percentage of spleen monocytes increases 4-fold in mice transplanted with aged HSCs, while those of NK cells, plasmacytoid DCs or mastocytes decrease up to 10-fold (Extended Data Fig. 12B- E).

Collectively, these results indicate that, similar to the fetal to postnatal transition, lymphoid architecture and B-cell differentiation pathways are subjected to a further developmental transition after the age of 60 years.

### Ontogeny-related variations in lymphoid production patterns are developmentally programmed

To search for age-dependent variations in lymphoid potential, HSCs or LMDPs (Extended Data Fig. 7 [Strategy-3]) from 26 donors between 17 PCW and 87 years were subsequently seeded under *scf:tpo:tslp* condition. FACS analysis of their differentiated progeny confirmed that HSC expansion rates decrease rapidly after birth and showed that, conversely to myeloid granulocyte or monocyte production that remains stable over age (data not shown), lymphoid outputs follow a downward trend, especially in donors over 50 years of age in whom high inter-individual and inter-clone differences are also observed (Fig. 7A). Similar age-dependent decline in lymphoid output is also seen in postnatal LMDPs after puberty which stresses the relevance of these findings (Fig. 7B). Most importantly, analyses of ELP production patterns confirmed that, in both HSC and LMDP compartments, transition to postnatal life is associated with a shift toward CD127^+^ ELPs whose relative percentages also gradually increase over age (Fig. 7C, D). Overall similar proliferation and differentiation patterns were observed in cultures under *scf:g:gm* condition, reinforcing the view that CD127^+^ ELPs are subjected to a cell-intrinsic regulation (Extended Data Fig. 11A-D). Consistent with these results, single-cell differentiation assays showed that postnatal HSPCs have decreased cloning efficiency, reduced expansion rates, and that they display predominant CD127^+^ ELP output (Extended Data Fig. 11E-L). Immunophenotypic characterization of *in vitro* differentiated ELPs confirmed that CD7 expression decreases as a function of donor age but failed to disclose concomitant variation in the expression pattern of CD10 (Fig. 7E), which argues for a more complex regulation of this marker. Lastly, culturing postnatal HSPCs under *scf:flt3l* condition revealed that, independent of the donor age, Flt3L fully restores the differentiation of the CD127^-^ ELPs (Fig. 7F). Collectively, these results show that ontogeny-related changes in the patterns of lymphocyte production are developmentally imprinted at the level of HSCs. In addition, they suggest that preservation of Flt3L responsiveness in adult donors could help maintain lymphoid homeostasis throughout postnatal life.

## DISCUSSION

In this study we followed the variations in lymphoid development architecture and lymphocyte production patterns taking place in 140 blood and BM donors between 13 development weeks and 87 years. As well as resolving longstanding controversies as to the phenotype and function of human MLPs, our data reveal that human lymphopoiesis undergoes three successive developmental transitions - the first at birth, then at puberty and lastly during aging - indicating that the developmental complexity of the lymphoid lineage has to date been largely underestimated.

Subjecting neonatal cord blood progenitors to optimized sorting schemes that efficiently resolve lymphoid lineage fates found that previously characterized CD127^-^ and CD127^+^ ELPs ^18^ circulate at low levels in umbilical cord blood and that, among them, differential CD7 versus CD10 expression distinguishes NK/ILC/T from B lineage-polarized subsets ^15, 29^. As expected, these analyses also confirmed that CD127^-^ and CD127^+^ ELPs correspond to the immediate downstream progeny of lymphoid-restricted CD117^lo^ MLPs (Alhaj Hussen et al. submitted). Subsequent functional characterization of neonatal MLPs revealed that differential CD7 and CD10 expression allows distinction between embryonic, fetal, and postnatal subsets and showed that, whereas embryonic CD7^+^ MLPs display balanced ELP production patterns, their fetal CD7^-^ and postnatal CD10^+^ counterparts are intrinsically biased toward CD127^+^ ELPs and the subsequent generation of B lymphocytes. Although our results are in line with earlier reports that independent subsets of CD7^+ 21, 37^ or CD10^+^ MLPs ^16, 17^ circulate in the neonatal blood, they show that the limited sensitivity of current multi-lineage differentiation assays has led to the erroneous conclusion that differential CD7 expression does not affect their lymphoid potential. Transcriptional profiling of neonatal MLPs confirmed their lymphoid affiliation but failed to disclose subset-specific gene signatures. Consistent with the idea that lymphocyte production patterns are conditioned at epigenetic level, ATAC-seq profiling found that neonatal CD10^+^ MLPs display increased accessibility of B-lineage genes.

Dynamic follow-up of lymphoid progenitors in the blood and BM of primary fetal and postnatal donors subsequently found that overall lymphoid outputs remain at stable levels until puberty after which they undergo a gradual age-dependent decline ^38^. Most importantly, these analyses revealed that a developmental shift from fetal multilineage to postnatal B lineage-biased lymphopoiesis takes place within the first few days after birth. Indeed, we found that transition to aerial life is associated with a global immunophenotypic CD7 to CD10 conversion of all lymphoid progenitor subsets and an increase in production of CD127^+^ ELPs which persists until puberty. In vitro and in vivo functional assays subsequently confirmed that these changes are developmentally imprinted and determined as early as the level of HSCs. Although the molecular bases of the neonatal shift in lymphocyte production patterns still need to be clarified, the temporal concordance with the fetal to adult switch in erythropoiesis strongly argues for involvement of conserved gene regulatory networks ^39–41^. Dynamic follow-up of age-related changes in lymphoid architecture also revealed a dual ontogeny and differentiation value of CD7 and CD10 markers. We found that while CD7 expression by fetal and neonatal HSCs reflects an early developmental origin, most probably during the first gestation trimester, its acquisition by downstream ELPs marks their specification toward the T/NK-ILC lineage ^15, 18^. An early embryonic origin of the CD7^+^ HSCs is further supported by transcriptional analyses showing that their gene signature is consistent with a more primitive developmental status.

However, so far it remains unclear whether the decreasing CD7 expression observed over age could also represent a surrogate marker of the decline in T-cell differentiation potential ^42^. On the other hand, the early acquisition of CD10 expression by lymphoid progenitors observed within the first few days birth can be considered as an immunophenotypic correlate of the B-lineage bias of postnatal lymphopoiesis.

Our results also provide evidence that the biological function and differentiation status of CD10^+^ MLPs change over age. Consistent with earlier reports ^32, 33^, the observation that CD10^+^ MLPs are enriched in the blood of pediatric donors indicates that, at least during the prepubertal period, this population displays a selective capacity to enter the blood flow and seed peripheral lymphoid organs, including thymus. This issue is even more important that CD10^+^ MLPs are detected in the thymus of pediatric donors where they are subjected to complex transcriptional rewiring prior entering the T-cell differentiation pathway ^43^. Our assumption that before puberty CD10^+^ MLPs have only a marginal contribution to BM lymphopoiesis is also supported by immunophenotypic analyses showing that, under steady state or regenerative conditions, CD127^+^ ELPs upregulate CD10 and CD19 as they progress along the B-lymphoid pathway. Immunophenotypic profiling of BM lymphoid progenitors from adult donors added further complexity to this picture by demonstrating that with increasing age percentages of CD10^+^ MLPs vary inversely with CD127^+^ ELPs. These observations point to a previously unknown developmental transition in the B-cell differentiation pathways with age in adults over 60 years. In contrast to younger donors, in which B-cell differentiation proceeds via CD127^+^ ELPs (whose highly proliferative status should also contribute to pre-commitment amplification of the B compartment), in elderly donors B-cell differentiation starts directly at the level of the CD10^+^ MLPs, bypassing CD127^+^ ELPs. Subsequent characterization of CD34^+^ HSPCs from immunodeficient mice reconstituted with HSCs from elderly donors confirmed that, in the same manner as the fetal to postnatal transition in lymphocyte production patterns, this late switch in the B-cell differentiation pathway is developmentally imprinted at the level of HSCs.

In conclusion, this study contributes to our understanding of the dynamics of lymphocyte production patterns during development and aging. Our results suggest that whereas fetal lymphopoiesis allows prenatal acquisition of diversified T and B cell repertoires essential to reach immunocompetence at birth, the neonatal boost in B cell production may be important to help broaden the antibody repertoire and enhance antibody-mediated immune responses to reduce morbidity and mortality due to infections during infancy. It remains unclear to date, whether the postnatal shift in lymphopoiesis also contributes to the decline in T cell production ^44^ or to age-related differences in the incidence of B- and T-cell acute lymphoblastic leukemia ^45^.

### AUTHOR CONTRIBUTIONS

S. K. & E. C. designed and performed most experiments; S. D. & F. J. analyzed the clonal data; S. D. & E. N. conducted the mouse studies; J. L. & T. D. provided the UCB; M. S., C. A. & F. G. provided the fetal and adult BM samples; A. C., E. V. contributed to data analyses and wrote the paper; D. G. & M. G. performed the ATAC-seq analyses; F. C., B. E. and T.A.D. performed the transcriptome analyses; K. A. H. and B. C. ensured the scientific supervision of the project and wrote the paper.

## ACKNOWLEDGEMENTS

The authors thank Christelle Doliger, Niclas Setterblad and Claire Maillard (Plateforme d’Imagerie et de Tri Cellulaire, IRSL, Paris France). We are grateful to Laurent David (Inserm 1064, Nantes) and Bernard Jost (GenomEast platform, IGBMC, Strasbourg, France). We also thank Jean Christophe Bories and Michele Souyri for critical discussions. This work was supported by the Agence de la Biomédecine, Agence National de la Recherche (ANR EpiDev), the Institut National du Cancer (InCA B-REC), the Fondation Ramsay Générale de Santé and by the INSERM HuDeCA network.

The authors have no conflict of interest.

## METHODS

### Human Sample Collection

Human fetal (gestational ages 13 to 23 weeks, n= 14) or postnatal (2-87 years, n=50) tissues from male or female donors were collected after informed consent according to institutional guidelines and the French bioethical legislation. Umbilical cord blood (UCB) was provided by the Unité de Thérapie Cellulaire of Hôpital Saint-Louis (Paris). Postnatal bone marrow (BM) samples were from healthy graft donors (Unité de Thérapie Cellulaire, Hôpital Saint-Louis, Paris). Fetal BMs were from spontaneously terminated pregnancies (Service de Biologie du Développement, Hôpital Robert Debré, Paris). Fetal (gestational ages 19 to 38 weeks, n= 13) and neonatal (0 to 77 days, n= 34) blood samples were from healthy donors (Service de Biologie du Développement, Hôpital Robert Debré, Paris).

### Mice

NOD.Cg-Prkdc*^scid^*IL2RG*^tm1wjl^*/SzJ (005557) mice known as NOD scid gamma (NSG) mice (Jackson Laboratory, Bar Harbor, MI) were housed in the pathogen-free animal facility of Institut de Recherche Saint Louis (Paris). Female NSG mice) were xenografted at 2 months of age. The Ethical Committee at Paris Nord University approved all performed experiments.

### Processing of human tissues and cell separation

Femurs from fetal or adult donors were sterilely excised and processed in RPMI 1640 medium supplemented with 10 U/ml RNase-free DNase I, before single-cell suspensions were filtered through a cell strainer (70 µm; BD biosciences). Blood or BM mononuclear cells were separated by Ficoll-Hypaque centrifugation (Pancoll, PAN Biotech GmbH) before processing for flow cytometry, cell sorting or CD34^+^ HSPC isolation. CD34^+^ HSPCs were isolated with the CD34 Microbead kit (Miltenyi Biotech; purity >90%), frozen in heat-inactivated fetal calf serum (FCS) supplemented with 10% DMSO and stored in liquid nitrogen until use.

### Xeno-transplantations

NOD scid gamma (NSG-SGM3) mice (Jackson Laboratory, Bar Harbor, MI) were housed in the pathogen-free animal facility of Institut de Recherche Saint Louis (Paris). The Ethical Committee at Paris Nord University approved all performed experiments. 3 months old mice were irradiated with 2.25 Gy 24 h before injection of 2.5 x 10^5^ CD34^+^ HSPCs in the caudal vein and sacrificed 8 weeks later. Single-cell suspensions recovered from femurs and tibias were filtered through a cell strainer (70 µm; BD biosciences) and depleted in mouse cells using the Mouse Cell Depletion Kit (Miltenyi) before being processed for Flow Cytometry.

### Flow cytometry and cell sorting

Human single-cell suspensions were incubated with human Fc receptor-binding inhibitor (Fc Block, eBioscience) before surface staining with anti-human monoclonal antibodies (mAbs). Fluorescence minus one (FMO) isotype controls were used to define positive signals for flow cytometry or cell sorting. Dead cells were excluded with the Zombie Violet Fixable Viability Kit (Biolegend). For labeling, cells were resuspended in PBS, 2 % FCS (1 to 5 x10^7^ cells/500 μl) and incubated with the following mAbs: CD45 AF700 (Biolegend, clone HI30), CD34 PB (Biolegend, clone 581), CD7 FITC (Beckman Coulter, clone 8H8.1), CD38 PerCPCy5.5 (Biolegend, clone HIT2), CD33 BV785 (Biolegend, clone WM53), CD115 APC (Biolegend, clone 9-4D2-1E4), CD71 PE (BD Bioscience, clone M-A712), ITGB7 PC7 (eBioscience, clone FIB504), CD3 PE-CF594 (BD Bioscience, clone UCHT1), CD14 PE-CF594 (BD Bioscience, clone MOP9), CD19 PE-DAZZLE (Biolegend, clone HIB19), CD24 PE-CF594 (BD Bioscience, clone ML5), CD56 PE-CF594 (BD Bioscience, clone B159), CD45RA BV711 (Biolegend, clone HI100), CD116 APC-vio770 (Miltenyi, clone REA211), CD127 PC5 (Biolegend, clone A019D5), CD117 BV605 (Biolegend, clone 104D2), CD10 BV650 (BD bioscience, clone HI10a), CD123 PE-Cy5.5 (Beckman Coulter, clone SSDCLY107D2), also CD45RA PE (BD Bioscience, clone HIT100), CD19 BV711 (BD bioscience, clone HIB19). Flow cytometry and cell sorting were performed with a BD Fortessa Analyzer or a BD FACSAria III sorter (BD Biosciences; (purity ≥ 95%). Flow cytometry analyses were performed using the FlowJo software (Version 10.7).

### Spectral cytometry

Human single-cell suspensions isolated from the spleen or BM of xenografted mice were processed and labeled as above were the following antibodies: CD45 PerCp (Biolegend, clone HI30), CD34 BUV615 (BD bioscience, clone 581), CD45RA BUV395 (Biolegend, clone HI100), CD38 APC-Fire 810 (Biolegend, clone HIT2), CD33 PE Texas Red (Biolegend, clone WM53), CD117 BV605 (Biolegend, clone 104D2), CD71 PECy5 (BD bioscience, clone M-A712), CD115 APC (Biolegend, clone 9-4D2-1E4), CD116 AP-Vio770 (Miltenyi, clone REA211), CD123 SB436 (BD bioscience, clone 7G3), CD127 APC R700 (Biolegend, clone A019D5), ITGB7 PECy7 (eBioscience, clone FIB504), CD7 FITC (Beckman Coulter, clone 8H8.1), CD10 BV650 (BD bioscience, clone HI10a), CD2 PerCpCy55 (Biolegend, clone RPA-2.10), CD19 Spark NIR 685 (Biolegend, clone HIB19), CD24 PE-AF 610 (Invitrogen, clone SN3), CD20 Pacific Orange (Invitrogen, clone HI47), IgM BV570 (Biolegend, clone MHM-88), IgD BV480 (Biolegend, clone IA6-2), CD5 BV785 (BD bioscience, clone UCHT2), CD56 BUV737 (BD bioscience, clone B159), CD94 PE (BD bioscience, clone HP-3D9), CD3 BUV661 (BD bioscience, clone HIT3a), CD1c AF647 (BD bioscience, clone F10/21A3), CD141 BB515 (BD bioscience, clone 1A4), CD303 BV750 (BD bioscience, clone V24-785), CD304 BV711 (Biolegend, clone 12C2), CD14 Spark Blue 550 (Biolegend, clone 63D3), CD16 BUV496 (BD bioscience, clone 3G8), CD15 BUV805 (BD bioscience, clone W6D3) and Zombie UV for dead cells. Spectral cytometry was performed with a Cytec Aurora cytometer (Spectro-Flo software). Flow cytometry analyses were performed using the FlowJo software (Version 10.7).

### Cell cultures and analysis

#### differentiation assays

For T cell differentiation assays, OP9-DL4 cells (1.5 x 10^3^ cells/well in 200 μl; gift of A. Cumano, Institut Pasteur, Paris) were seeded in Opti-MEM-Glutamax supplemented with 10% FCS, 1% penicillin-streptomycin and 1/1000 *β*-mercapto-ethanol (Life Technologies) in 96-well flat-bottom plates 24 hour prior to co-culture. Before cell sorting, medium was removed and 200 μl of complete medium supplemented with SCF (10 ng/ml), FLT3L (5 ng/ml), and IL-7 (2 ng/ml) (all from Miltenyi Biotech) was added. 50 Cells were sorted directly into the wells and allowed to differentiate for 2 weeks. On culture day 7, 50 μl of complete medium supplemented with 4X cytokine concentrations were added to each well. At culture day 14, the well content was harvested, and cells were labeled with CD45 AF700 (Biolegend, clone HI30), CD34 PB (Biolegend, clone 581), CD7 BV510 (BD Bioscience, clone M-T701), CD5 APC (BD Bioscience, clone UCHT2), CD4 FITC (Biolegend, clone RPA- T4), and CD8a PC7 (Biolegend, clone RPA-T8) mAbs.

For NK differentiation assays, OP9 cells were seeded as above in 96-well flat-bottom plates 24 hour prior to culture initiation. Before cell sorting, medium was removed and 200 μl of complete medium supplemented with SCF (10 ng/ml), FLT3L (10 ng/ml), IL-7 (10 ng/ml) and IL-15 (10 ng/ml) (all from Miltenyi Biotech) was added. Cells were sorted directly into the wells (50 cell/well) and allowed to differentiate for 2 weeks. On culture days 7, 50 μl of fresh medium supplemented with 4X cytokine concentrations were added to each well. At culture day 14, the well content was harvested, and cell surface staining was performed with the following mAbs: CD45 AF700 (Biolegend, clone HI30), CD7 FITC (Beckman Coulter, clone 8H8.1), CD56 PC7 (Biolegend, clone MEM-188), CD14 PE-CF594 (BD Bioscience, clone MOP9), CD94 APC (Biolegend, clone DX22). NK cells were defined as the CD45^+^CD14^-^CD56^+^ subset.

For B cell differentiation assays, MS5 cells (1.5 x 10^4^ cells/well) were seeded 24 hours prior to co-culture initiation in 96-well flat-bottom plates with RPMI medium supplemented with 10% FCS, 1% glutamine, and 1% penicillin-streptomycin and 1/1000 *β*-mercapto-ethanol (Life Technologies).

Before cell sorting, medium was removed and 200 ml of complete medium supplemented with SCF (10 ng/ml), TPO (10 ng/ml), and IL-7 (10 ng/ml). Cells were sorted directly into the wells by pools of 50 to 100 cells and allowed to differentiate for 2 weeks. On culture day 7, 50 ml of complete medium supplemented with 4X growth factor concentrations were added to each well. The following antibodies were used to assess BL differentiation: CD45 AF700 (Biolegend, clone HI30), CD19 APC (Biolegend, clone HIB19), CD9 PE-CF594 (BD biosciences clone M-L13), CD7 FITC (Beckman Coulter, clone 8H8.1), CD10 BV650 (BD bioscience, clone HI10a), CD20 APC (Biolegend, clone 2H7), CD24 BV785 (BD biosciences clone ML5), CD127 PC5 (Biolegend, clone A019D5), IgM BV421 (Biolegend, clone MHM-88), IgD PC7 (Biolegend, clone IA6-2). Flow cytometry was performed as above with a BD Fortessa Analyzer and the FlowJo software.

#### diversification assays

For diversification assays under bulk (100 cell/well) or clonal conditions, OP9 stromal cells were seeded as above in Opti-MEM-Glutamax supplemented with 2.5% FCS, 7.5% BIT 9500 (StemCell Technologies), 1% penicillin-streptomycin and 1/1000 b-mercapto-ethanol (Life Technologies) in 96- well U-bottom plates 24 hours prior to co-culture with the indicated combinations of growth/differentiation factors (10 ng/ml each; all from Miltenyi Biotec). UCB or BM CD34^+^ HSPC subsets were directly seeded by FACS in the corresponding plates and cultured for 10 (bulk cultures) or 14 (clonal cultures) days before being subjected to FACS analysis. Flow cytometry analyses were performed with a BD Fortessa Analyzer using the following antibodies: CD45 AF700 (Biolegend, clone HI30), CD34 PB (Biolegend, clone 581), CD7 FITC (Beckman Coulter, clone 8H8.1), CD123 BV786 (BD Bioscience, clone 7G3), CD115 APC (Biolegend, clone 9-4D2-1E4), CD45RA PE (BD Bioscience, clone HIT100), ITGB7 PC7 (eBioscience, clone FIB504), CD24 PE-CF594 (BD Bioscience, clone ML5), CD19 BV711 (BD bioscience, clone HIB19), CD116 APC-vio770 (Miltenyi, clone REA211), CD127 PC5 (Biolegend, clone A019D5), CD15 BV605 (BD bioscience, clone HI10a), CD10 BV650 (BD bioscience, clone HI98). Flow cytometry analyses were performed using the FlowJo software.

### Chromatin accessibility mapping by ATAC-seq

Transposing reactions for ATAC-seq were performed as previously described ^46^, using between 15- 30 000 cells from the indicated populations (minimum 2 biological replicates per population). Sequencing libraries were generated using 5 cycles of PCR pre-amplification with Nextera barcoding primers ^46^ followed by 3 to 5 additional cycles, according to the quantity of pre-amplified product as detected in a qPCR side reaction ^47^. Amplified libraries were size selected and sequenced (100 bases paired end) on a HiSeq 4000 to a minimum depth of 30 million raw paired reads, at the GenomeEast Platform of the IGBMC, Strasbourg, France.

### Gene expression profiling by mini-RNA-seq

The protocol used for mini-RNAseq was adapted from Soumillon et al. ^48^. For each population batches of 100 cells (2-10 replicates) were sorted by FACS in 96-well V-bottom plate wells containing 2 μl of lysis buffer (Ultra-pure Water containing 10% Triton X-100 and RNasin plus 40U/ul). After evaporation of the lysis buffer at 95°C for 3 minutes, first strand cDNA synthesis was performed using Maxima H Minus Reverse Transcriptase (Thermo Fisher), E3V6NEXT primers (specific primer for each well) and Template Switching Primer: E5V6NEXT. Resulting cDNAs were then pooled and purified by DNA Clean and Concentrator TM-5 (Zymo research) and treated with Exonuclease I (New EnglandBiolabs) before amplification by the Advantage 2 PCR kit (Clontech) and the SINGV6 primer (95°C for 1 min, 15 cycles at 95°C for 15s, 65°C for 30s, 68°C for 6 min, and 72°C for 10 min). PCR products were purified by Agencourt AMPure XP (Beckman Coulter). Libraries were prepared using Illumina DNA Prep (Illumina) according to the manufacturer’s guidelines and sequenced on a HiSeq 4000 at the GenomeEast Platform of the IGBMC (Strasbourg, France). Image analysis and base calling were performed using RTA 2.7.7 and bcl2fastq 2.17.1.14. Adapter dimer reads were removed using DimerRemover (https://sourceforge.net/projects/dimerremover/).

## Quantification and statistical analyses

### Algorithm-assisted clone classification

Immunophenotypic data were extracted using the FlowJo software following the gating strategy described in Extended Data Fig. 3A. Algorithm-assisted clone classification was based on size (absolute cell number/clone) and relative percentages of HSPCs (CD34^+^), granulocytes (CD15^+^), monocytes (CD115^+^), dendritic cells (CD123^+^) and lymphoid cells. Percentages of lymphoid cells per clone were defined as the sum of the percentages of CD19^+^ B cells, CD127^-^ ELPs and CD127^+^ ELPs. Data were processed with the ClonAn software (see below) using the UMAP package (http://arxiv.org/abs/1802.03426) ^49^. The ClonAn software used for algorithm-assisted clone classification analyses was developed by Samuel Diop on top of the Scikit-Learn Library ^50^.

### ATAC-seq analysis and bioinformatics

Demultiplexed sequencing data were processed using the ENCODE ATAC-seq pipeline version 1.7.2 (https://github.com/ENCODE-DCC/atac-seq-pipeline) using the hg38 assembly and the default parameters for paired end sequencing. Only samples with TSS enrichment >5 were used in subsequent analyses. Downstream analyses were performed using the optimal peakset output (FDR<0.05) from Irreproducible Discovery Rate analysis. Peaks within 500 bp were merged using bedtools MergeBED (-d 500) ^51^. Reads in peaks were counted using Featurecounts ^52^ with default parameters for paired end reads. PCA of signal within peaks and detection of differential peaks (log2 fold change>0.5; padj<0.05) were carried out using EdgeR with TMM normalisation and robust conditions ^53^. Coverage bigwig files were generated with bamCoverage (deepTools2) ^54^ using TMM-normalisation factors and were visualised on the UCSC genome browser together with Encode ChIP-seq tracks of H3K4me1 and H3K27ac in K562 and GM12878 (“https://www.encodeproject.org“). Normalised bigwig files were used to generate heatmap and pileup profiles using computeMatrix (scaled region mode; 1 kb flanking each peak, sorting on mean peak signal) followed by plotHeatmap (deepTools2). Distribution of peaks relative to genomic features (hg38) was analysed with ChIPseeker at default settings except with promoters defined as +- 500 bp of the TSS ^55^. Ontology analysis of genes associated with ATAC peaks was performed with the GREAT tool (v4.0.4) (3http://great.stanford.edu/public/html/“) ^56^, using the whole hg38 genome as background and applying association settings of Basal plus extension, proximal windows of 2 kb upstream and 1 kb downstream, plus distal up to 1 Mb and including curated regulatory domains. Motif analysis was performed using the XSTREME tool (v5.4.1) (Charles E. Grant and Timothy L. Bailey, “XSTREME: comprehensive motif analysis of biological sequence datasets”, *BioRxiv*, 2021). of the MEME suite^57^ at default parameters, using background sets of unchanging peaks of equivalent sizes to the test peaksets.

### Mini-RNA-seq analysis and bioinformatics

Data quality control and pre-processing were performed by SciLicium (Rennes, France). Briefly The first read contains 16 bases that must have a quality score higher than 10. The first 6 bp correspond to a unique sample-specific barcode and the following 10 bp to a unique molecular identifier (UMI). The second reads were aligned to the human reference transcriptome from the UCSC website using BWA version 0.7.4.4 with the parameter “−l 24”. Reads mapping to several positions in the genome were filtered out from the analysis. The complete pipeline has been previously described in ^58^. After quality control and data pre-processing, a gene count matrix was generated by counting the number of unique UMIs associated with each gene (lines) for each sample (columns). The UMI matrix was further normalized with the regularized log (rlog) transformation package implemented in the DeSeq2 package ^59^. Raw and pre-processed data are accessible on Gene Expression Omnibus (GEO): GSE215045; GSE215052.

The statistical comparisons were performed in the AMEN suite of tools ^60^. For each transcriptomic analysis, several relevant comparisons were selected to identify differentially expressed genes (DEGs). Briefly, genes showing an expression signal higher than 0.0 and at least a 2.0-fold-change between the two experimental conditions of each pairwise comparison were selected. The empirical moderated t-statistics implemented into the LIMMA package (F-value adjusted using the Benjamini & Hochberg (BH) False Discovery Rate approach, p ≤ 0.05) ^61, 62^ was used to define the set of genes showing significant statistical changes across all comparisons of a given transcriptomic analysis. The resulting set of DEGs were then clustered into distinct expression clusters with the k-means algorithm and presented as heatmaps by using the pheatmap R package developed by R. Kolde (2019) (version 1.0.12. “https://CRAN.R-project.org/package=pheatmap“).

Gene Ontology enrichment analysis was performed with the AMEN suite ^60^. A specific annotation term was considered enriched in a gene cluster when the FDR-adjusted p value was ≤0.05 (Fisher’s exact probability). The regulatory network representations were drawn using the igraph package (“https://igraph.org“) (Csardi & Nepusz, 2006) implemented in R. The regulation data described correspond to a consolidation of all gene sets provided by GSEA ^63^ associated with transcription factor targets. Those transcription factor target prediction gene sets are based on the Gene Transcription Regulation Database (GTRD) ^64^ and legacy sets, the latter corresponding to upstream cis-regulatory motifs ^65^.

## SUPPLEMENTARY TABLES

Table S1. Ontology analysis of chromatin changes in MLP 10p relative to HSC. Complete lists of Gene Ontology (GO) Biological Process categories enriched (FDR q<0.05) in genes associated with peaks of increased (left) or decreased (right) accessibility in 10p MLPs relative to HSCs. Shown for each category is -log10 of the binomial FDR q value for significance of enrichment. The association rule used for assigning peaks to genes in GREAT was basal+extension: 5000 bp upstream, 1000 bp downstream, 200000 bp max extension, curated regulatory domains.

Table S2. Gene expression profiling of 11 neonatal HSPC subsets. (A) Distribution of 1, 773 differentially expressed genes (DEGs) across 10 clusters (C1-10; fold-change ≥ 1.5; adjusted F-value

≤ 0.005); population-specific gene set enrichments are highlighted in gray. (B) Complete list of GO and GSEA categories enriched in clusters C1-10 (enrichment threshold: adjusted *p* value ≤ 0.001).

Table S3. Direct side-by-side comparison between fetal and postnatal BM CD34^+^ HSPCs. (A) Distribution of 1, 034 differentially expressed genes (DEGs) upregulated (n=511) or downregulated in fetal HSPCs (n=523) (fold-change ≥ 1.5; adjusted F-value ≤ 0.005); population-specific gene set enrichments are highlighted in gray. (B) List of the top 200 GO biological process categories enriched in fetal or postnatal HSPCs (adjusted p values are indicated).

Table S4. Gene expression profiling of 8 fetal HSPC subsets isolated from the BM of 2 donors (19-20 PCW). (A) Distribution of 517 differentially expressed genes (DEGs) across 8 clusters (P1-8; fold-change ≥ 1.5; adjusted F-value ≤ 0.005); population-specific gene set enrichments are highlighted in gray. (B) Complete list of GO and GSEA categories enriched in clusters P1-8 (enrichment threshold: adjusted *p* value ≤ 0.01).

Table S5. Gene expression profiling of 6 postnatal HSPC subsets isolated from the BM of 4 postnatal donors (11-52 years). (A) Distribution of 339 differentially expressed genes (DEGs) across 7 clusters (Q1-7; fold-change ≥ 1.5; adjusted F-value ≤ 0.005); population-specific gene set enrichments are highlighted in gray. (B) Complete list of GO and GSEA categories enriched in clusters Q1-7 (enrichment threshold: adjusted p value ≤ 0.01).

---

**Extended Data Figure 1.**
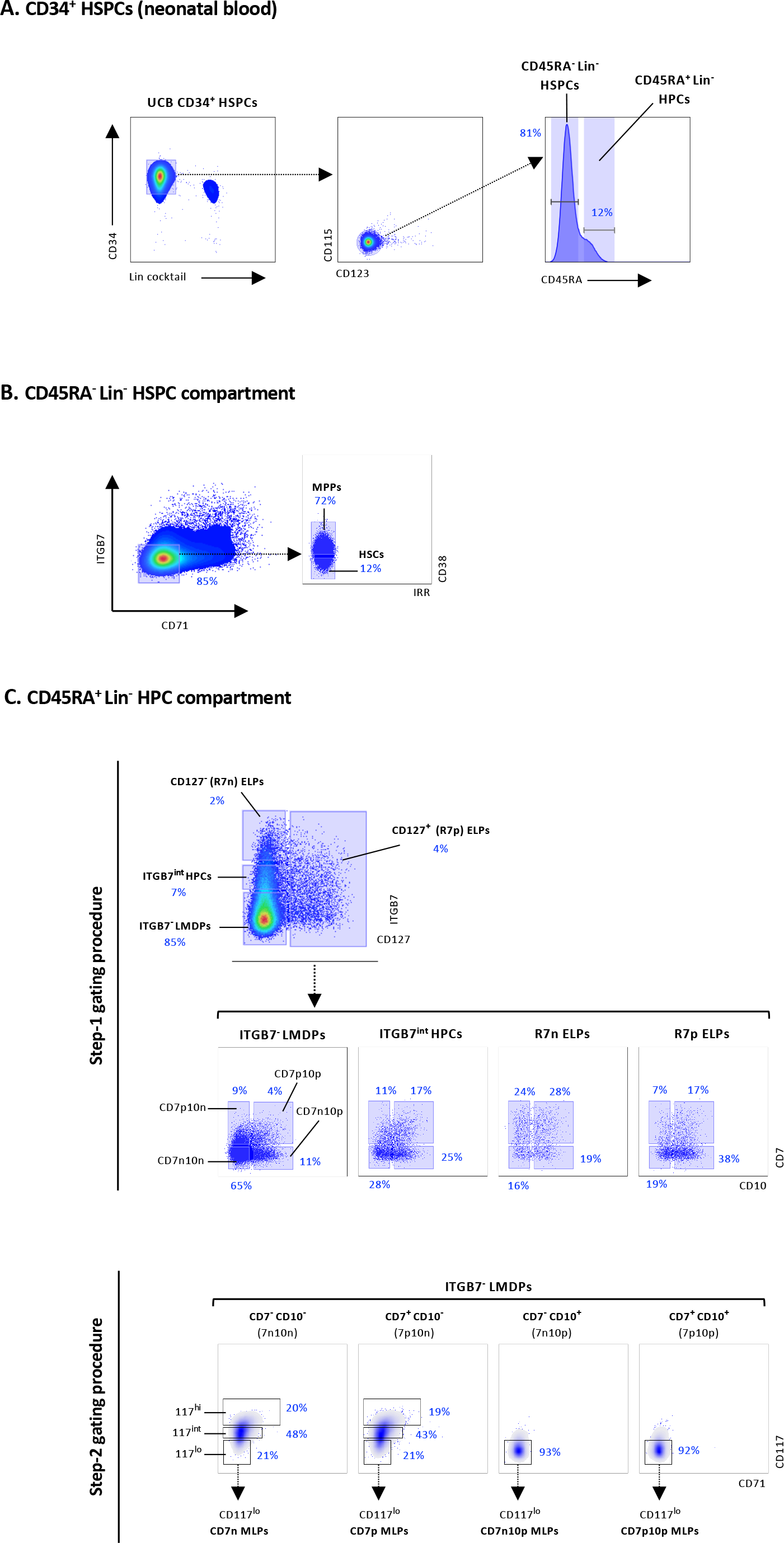
Detailed gating schemes used for molecular and functional characterization of neonatal HSPCs. Immunophenotypic procedure to suddivide neonatal HSPCs. CD34^+^ HSPCs freshly isolated from neonatal UCB were labeled with a 16 antibody panel and fractionated as follows. (A) Right pseudocolor plot shows selection of CD34^+^Lin^-^ HSPCs (Lin cocktail: CD3, CD19, CD14, CD24, CD56). After exclusion of CD115^+^ or CD123^+^ monocyte/dendritic precursors (My^-^), remaining cells were subdivided based on CD45RA expression into CD45RA^-^ or CD45RA^+^ compartment. (B) To isolate HSCs, CD45RA^-^Lin^-^ were further subdivided based on CD71 and ITGB7 expression to exclude erythro-megakaryocytic precursors (*Notta al. Science 2016, 351 (6269):aab2116*) before fractionation into distinct CD38^lo/-^ HSC or CD38^+^ MPP subsets (right panel). (C) Subdividivision of CD45RA^+^Lin^-^ HPCs based on differential expression of ITGB7, CD127 and CD7 or CD10 markers. Upper pseudocolor plot shows the subdivision of CD45RA^+^Lin^-^ cells into LMDP (ITGB7^-^), HPC (ITGB7^int^) or ELP (CD127^-/^CD127^+^) ELP subsets used for further fractionation procedures based on differential CD7/CD10 (Step-1) or CD117 (Step-2) expression.. <u>Step-1 fractionation procedure</u> (upper dot plots): each LMDP, HPC or CD127^-/+^ compartment was partitioned into CD7^-^CD10^-^ (7n10n), CD7^-^ CD10^+^ (7n10p), CD7^+^CD10^-^ (7p10n) or CD7^+^CD10^+^ (7p10p) subfractions; the resulting 16 cellular subsets plus CD45RA^-^CD38^lo/-^ HSCs, were subsequently cultured for 14 days (100 cell/well) under mono-lineage T, B or NK conditions; alternatively, the CD7np/CD10np LMDP subfractions were seeded in bulk or clonal diversification conditions or processed molecular profiling. <u>. Step-2 fractionation procedure</u> (lower density plots): the indicated CD7^-/+^CD10^-/+^ LMDP subsets were further subdivided into CD117^hi^ (20%), CD117^int^ (40%) or CD117^lo^ (20%) fractions before seeding in diversification assays or processing for mini-RNAseq; dashed arrows show the gating procedure. Percentages are indicated; data are from one representative experiment out of > 50.

**Extended Data Figure 2.**
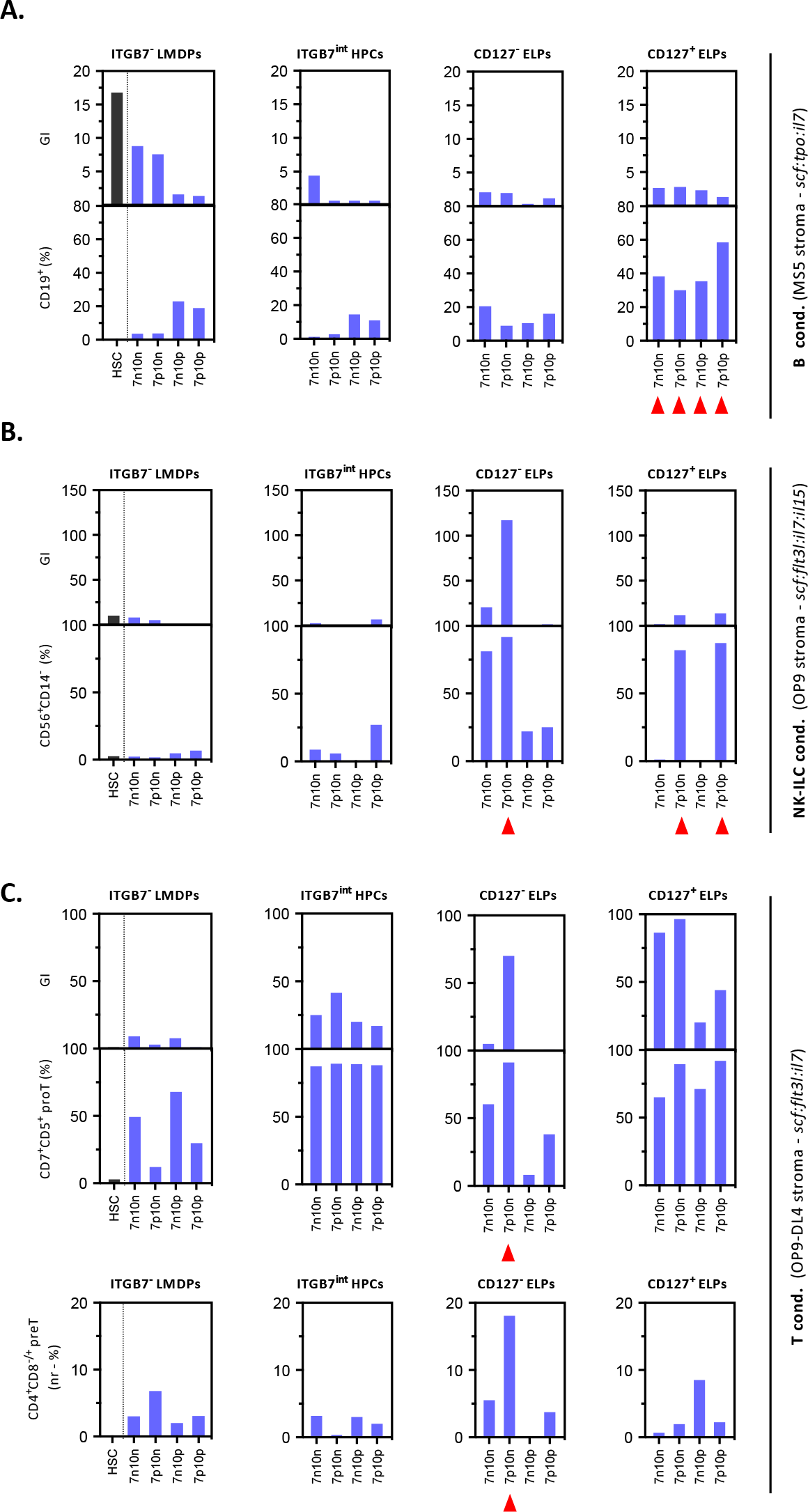
In vitro assessment of lymphoid potential in B, NK or T differentiation assays. The indicated cellular fractions (Extended data Fig. S1B: Step-1 gating procedure) were seeded by FACS in 96-well plates (100 cells/well) and cultured for 2 weeks under: (A) B (MS5 cells; *scf:tpo:il7*), (B) NK-ILC (OP9 cells; *scf:flt3l:il7:il15*) or (C) T (OP9-DL4 cells; *scf:flt3l:il7*) conditions before assessment of lineage outputs. Results are expressed as normalized growth indexes (GI) or percentages of in vitro-generated B (CD19^+^), NK-ILC (CD56^+^CD14^-^) or pro-T (CD7^+^CD5^+^) cells. For the T condition, percentages of pre- or post-β-selection CD4^+^CD8^-/+^ preT cells amongst CD7^+^CD5^+^ pro-T cells are also shown (lower panels). Blue bars correspond to median values of 10-40 replicates; results are pooled from ≥ 2 independent experiments for each cellular fraction and culture condition; red arrows indicate hotspots of lymphoid potential. Notably, in cultures under the T condition only CD127^-^CD7^+^ ETPs progressed through the β-selection checkpoint within a 2 week-period. It has also to be noted that within more immature LMDPs or HPCs, only the CD10^+^ cellular fractions had detectable B-cell potential.

**Extended Data Figure 3.**
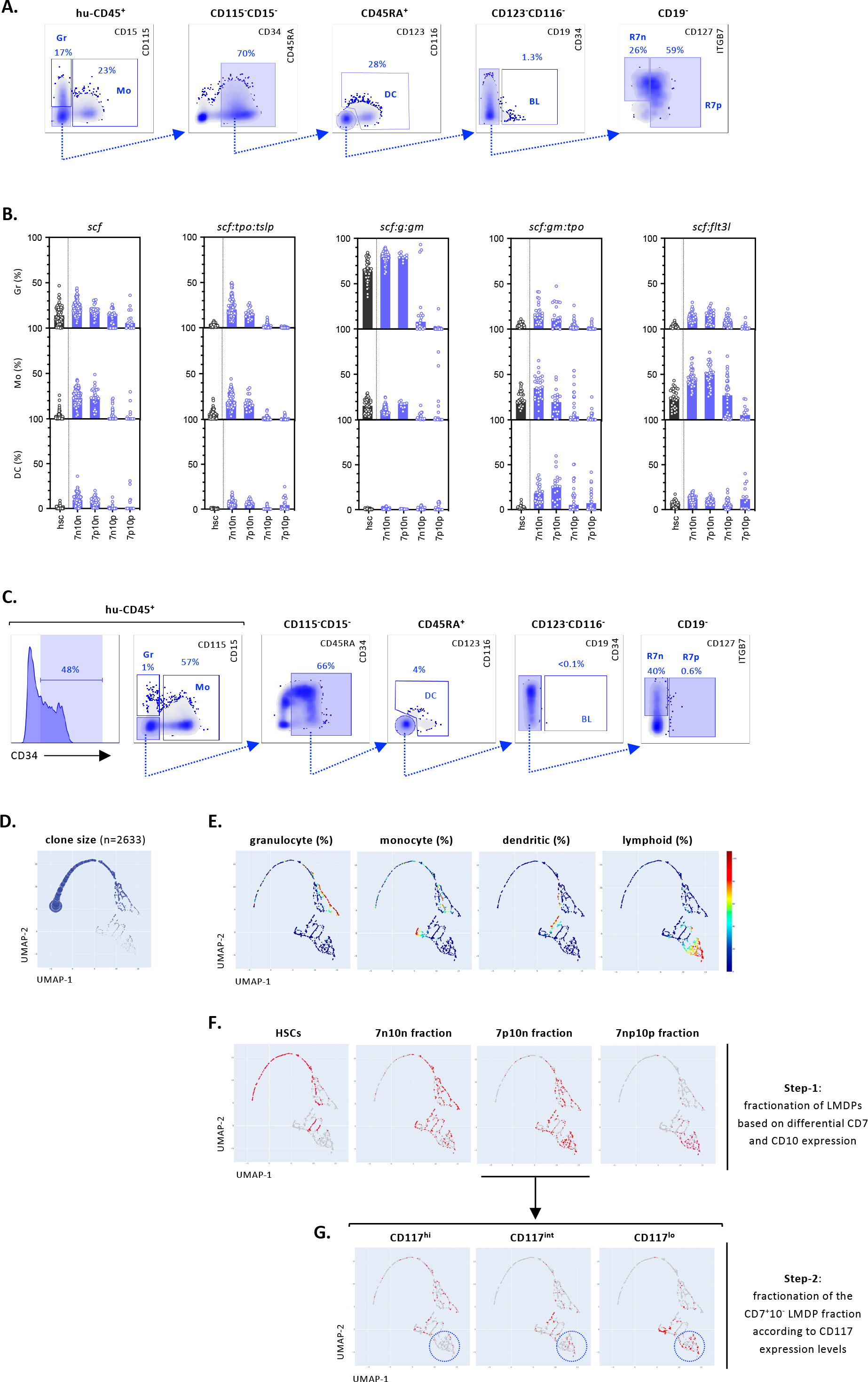
Assessment of lympho-myeloid differentiation potentials in bulk or clonal in vitro diversification assays. (A). Gating strategy for analysis of bulk diversification assays. LMDPs seeded by 100 cell-pools in 96-well culture plates were cultured for 10 days onto OP9 cells under standard *scf* condition before FACS analysis with a 13-antibody panel. After exclusion of doublets, gates were set on hu-CD45^+^ cells which were first divided into CD115^-^CD15^+^ granulocytic (Gr) or CD115^+^CD15^-^ monocytic (Mo) subsets. To capture dendritic cell precursors CD45RA^+^ (CD115^-^CD15^-^) cells were further subdivided based on CD116 and CD123, with the CD123^+^CD116^-/+^ subset being assigned to the dendritic cell lineage (DC). Remaining cells were further fractionated into CD19^+^ (BLs) or CD34^+^CD19^-^ subsets. Further fractionation of CD19^-^ cells according to ITGB7 and CD127 expression allowed isolation of CD127^-^ (R7^-^) or CD127^+^ (R7^+^) ELPs. Blue arrows indicate the gating procedure; bi-dimensional density and dot plots are from 10 concatenated wells; shown is expression of the indicated markers; percentages of the corresponding populations are indicated. (B). Quantification of myeloid granulocyte (upper panels), monocyte (medium panels) or dendritic (lower panels) outputs in diversification assays under *scf*, *scf:g:gm*, *scf:gm:tpo or scf:flt3l* condition. LMDPs fractionated based on differential CD7 and CD10 expression were cultured and analyzed as indicated above; bars show median percentages of the indicated subsets; empty circles correspond to individual wells. HSCs (black bars and circles)are used as internal controls. (C). Gating strategy for analysis of clonal diversification assays. HSC or LMDP subsets were seeded individually in 96-well plates and cultured for 14 days onto OP9 stroma under the *scf:gm-csf:tpo* condition before FACS analysis. After exclusion of doublets gates were set on hu-CD45^+^ cells. Left histogram and density plots show the strategy used for quantification of CD34^+^ HPCs, granulocyte (CD15^+^CD115^-^), monocyte (CD115^+^CD15^-/+^), or dendritic (CD123^+^CD116^-/+^) outputs; percentages of lymphoid cells are defined as the sum of CD19^+^ BL and CD127^-/+^ ELP percentages. Results are from a single HSC-derived multipotent clone; percentages of the corresponding populations are indicated. (D-G). Clonal diversification assays. Uniform Manifold Approximation and Projection (UMAP) algorithm was applied to the clonal data pooled from all experiments and ancestor cell types (n=2633). (D) Left bubble map shows distribution of ancestor cells according to clone size; (E) heatmaps show the distribution of granulocyte, monocyte, dendritic or lymphoid potentials defined as percentages of corresponding subsets within individual clones; snapshots show projection onto the UMAP of clones derived from (F) HSC or LMDP ancestors or (G) CD7^+^CD10^-^ (7p10n) LMDPs subdivided according to CD117^hi-int-lo^ expression; dashed circles show the area of projection of lymphoid clones.

**Extended Data Figure 4.**
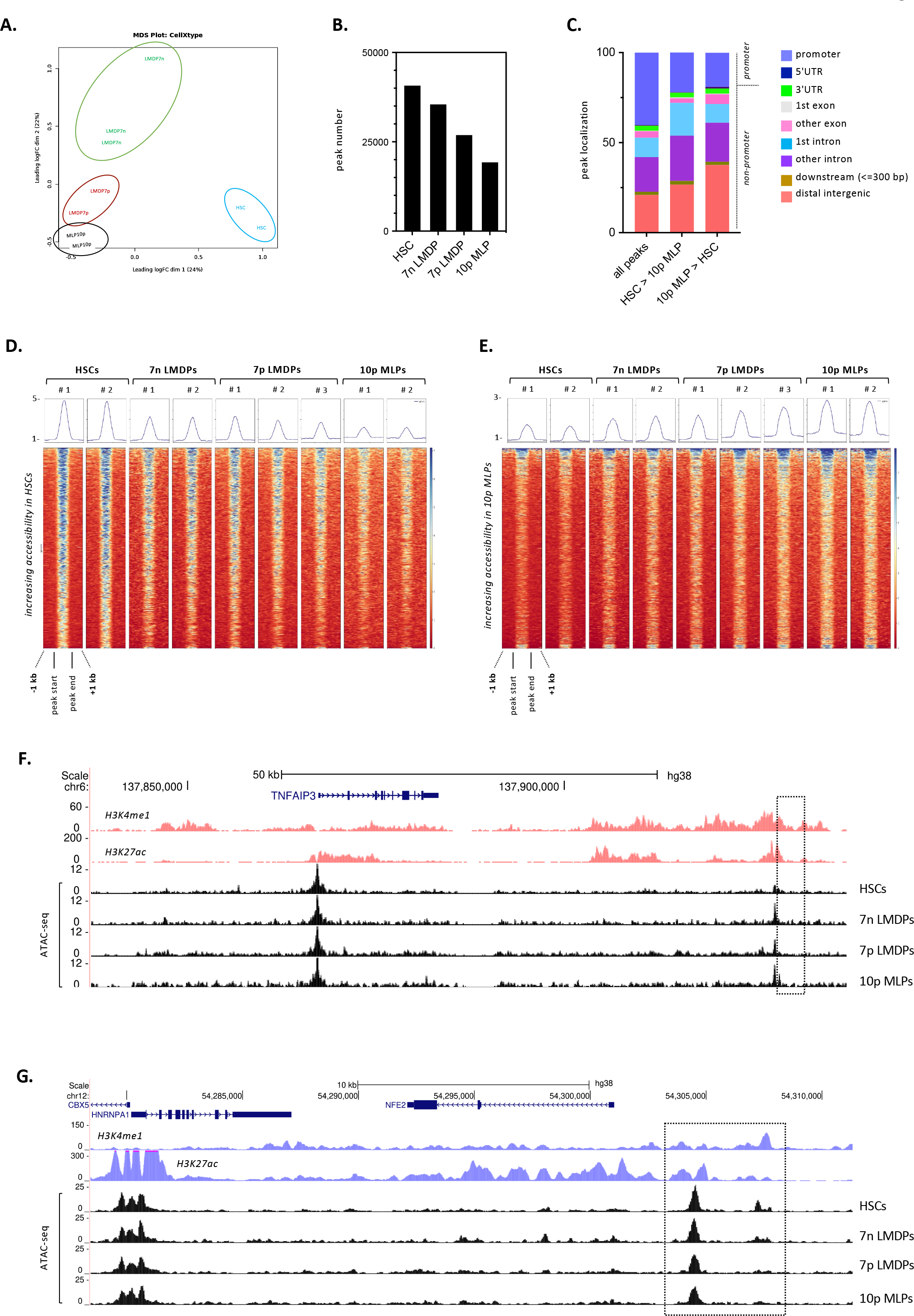
Chromatin accessibility landcape of neonatal HSPC subsets. ATA-seq profiling of neonatal CD34^+^ HSPCs. (A) MultiDimensional Scaling (MDS) plot from EdgeR indicating relatedness of samples based on normalized ATAC-signal at the master set of 46531 peaks (IDR FDR<0.05) observed in at least one population. (B) Total number of ATAC-seq peaks (IDR FDR<0.05) detected in the indicated populations and (C) distribution of indicated peak sets relative to genomic features. (D,E) Heat maps of normalised ATAC-seq signal in the samples indicated at 1125 peaks decreasing (D) or 1579 peaks increasing (E) in 10p MLPs relative to HSCs; a pileup representation of cumulative ATAC-seq signal is shown above each heatmap. (F, G) UCSC Genome Browser snapshots showing ATAC-seq signal in the indicated populations at two example genetic loci: (F) TNFAIP3 (from GO category : Immune response-activating signal transduction); (G) NFE2 (GO category : Regulation of myeloid cell differentiation). Gene annotation (Gencode V38) is shown at top followed by Encode tracks demonstrating enhancer-associated histone modifications (H3K4me1 and H3K27ac) in the K562 erythro-myeloid or GM12878 lymphoblastoid cell lines; regions whose accessibility increases during lymphoid specification are highlighted (dashed line).

**Extended Data Figure 5.**
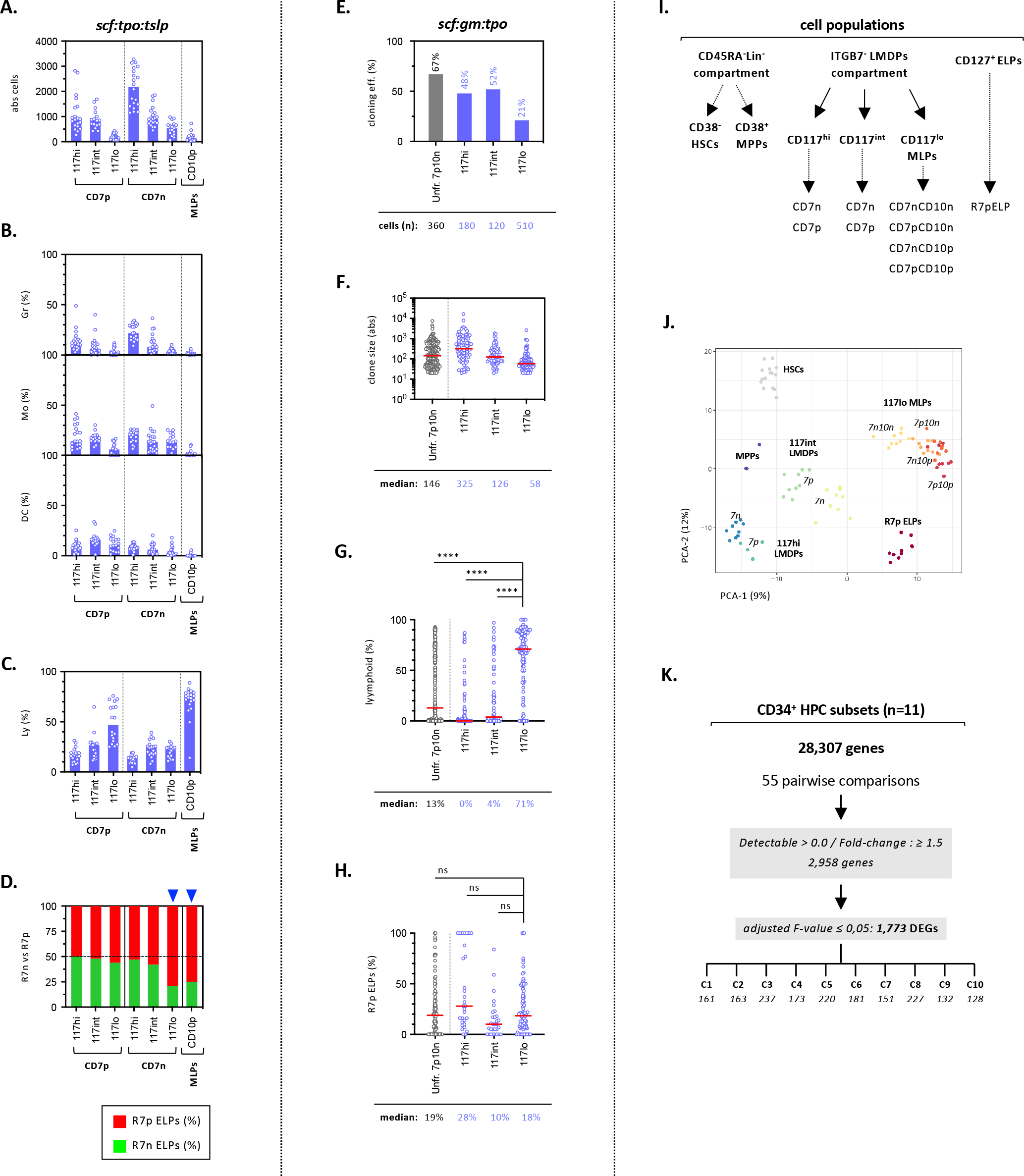
Molecular and functional characterization of neonatal MLPs. (A-D). Bulk diversification assays. The indicated CD7^-^(CD10^-^) or CD7^+^ (CD10^-^) LMDP subsets were subdivided into CD117^hi/int/lo^ fractions (Extended Data Fig. 1C : Step-2 gating procedure) and seeded for 10 days by 100-cell pools under the *scf:tslp:tpo* condition before quantification of cell growth and lineage output. Bar plots show (A) absolute cell numbers or percentages of (B) myeloid or (C) lymphoid cells; bars indicate medians; circles correspond to individual wells; assessment of statistical significance was performed with the Kruskal-Wallis test (****p <0.0001 for all conditions). (D) Stacked bar plots show the relative proportions of CD127^-^ (green) and CD127^+^ (red) ELPs; results are expressed as median percentages from ≥10 individual wells; blue arrows show statistically significant differences (CD117^lo^CD7n10n or CD10p versus others; p < 0.0001 by the Mann-Whitney test). (E-H). Clonal diversification assays. CD7^+^ LMDPs fractioned as above based on CD117 expression levels were seeded individually in 96-well plates and cultured for 14 days onto OP9 stroma before quantification of lineage outputs. Bar and circle plots show: (E) cloning efficiencies; (F) absolute cell number per clone (clone size); percentages of (G) total lymphoid cells (H) CD127^+^ ELPs per clone. CD127^+^ ELP percentages are normalized relative to total ELPs. For circles plots, red bars correspond to the median values shown below each plot; assessment of statistical significance was performed with the Mann-Whitney test (**** p<0.0001; *** p<0.001; ** p<0.01;). Results are pooled from 3 experiments. (I-K). Transcriptional profiling of neonatal HSPCs by mibi-RNA sequencing. (I) Experimental design: the indicated lineage negative (Lin^-^) CD45RA^-^ or CD45RA^+^ fractions were sorted from freshly isolated UCB CD34^+^ HSPCs, cultured for 3 hrs in cytokine-supplemented medium (*scf:gm:tpo*) and sorted again by 100 cell batches (3-9 replicates/population) before being processed for mini-RNAseq; (J) Principal component analyses (PCA) of transcriptome data; (K) Pipeline for detection of differentially expressed genes (DEGs); detection thresholds, fold-changes and adjusted F-values are indicated.

**Extended Data Figure 6.**
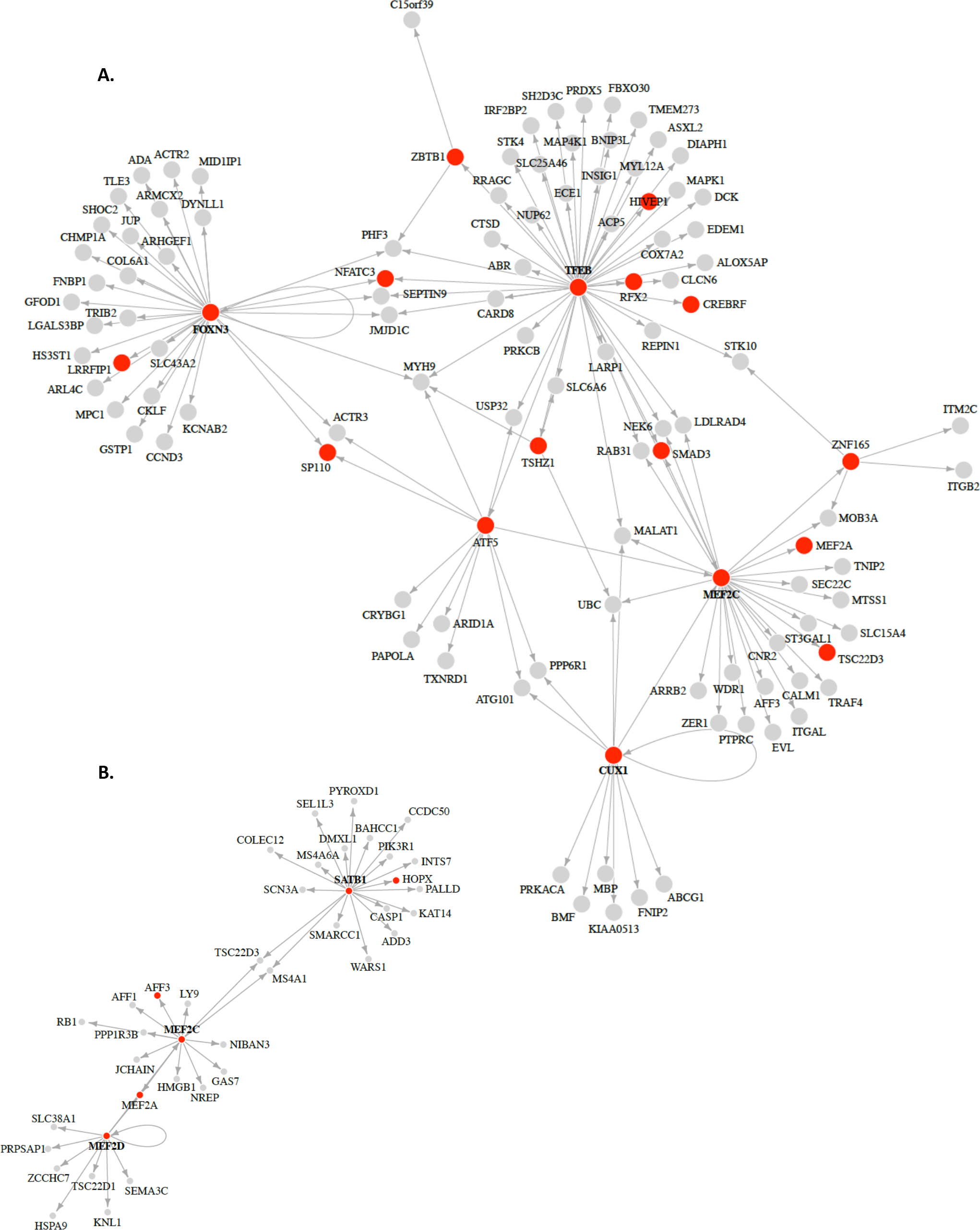
Transcription factor (TF) activity associated with lymphoid specification and prediction of downstream target genes. Connectivity plots of TFs (red dots) overexpressed in (A) UCB or (B) BM MLPs.

**Extended Data Figure 7.**
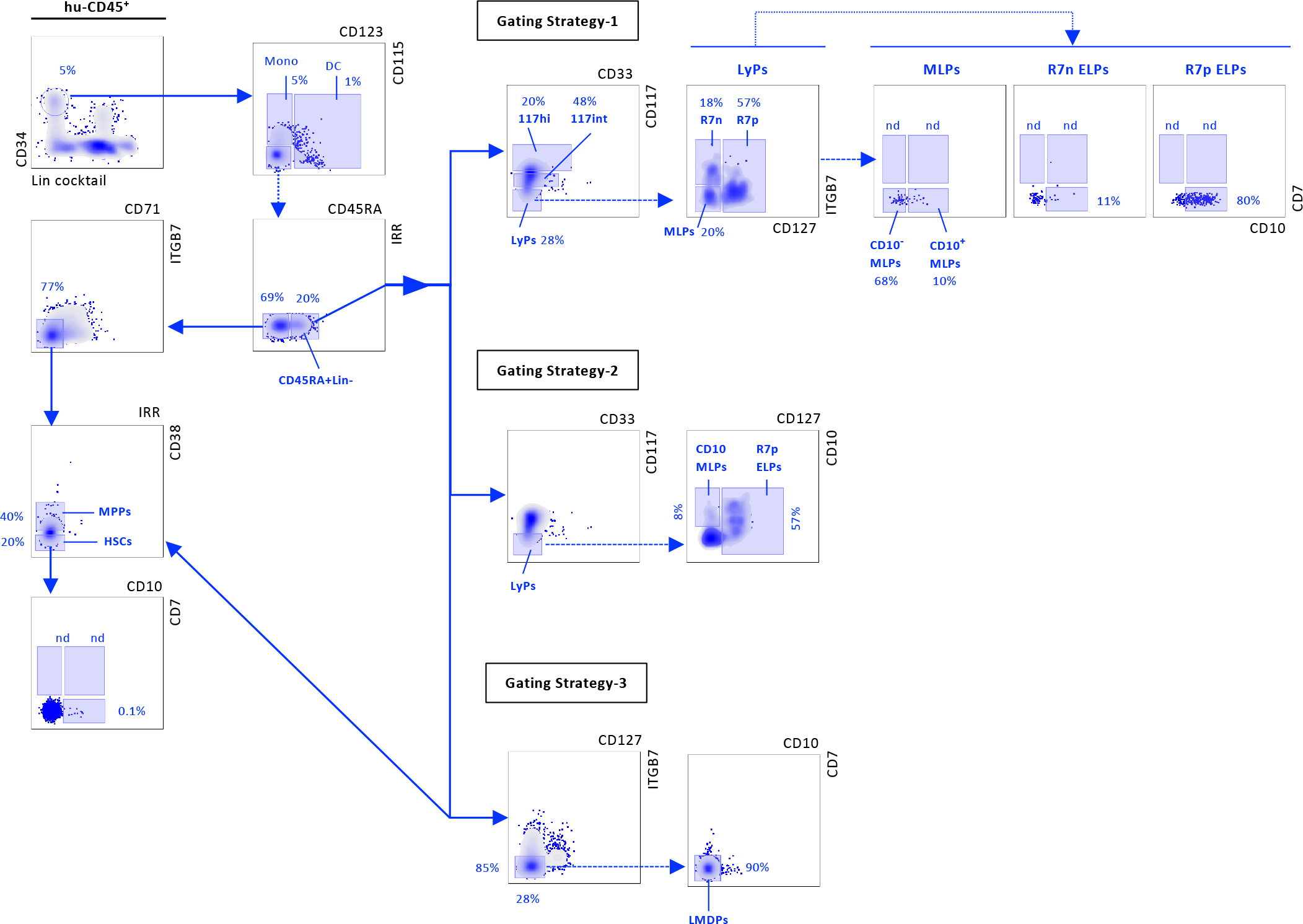
Detailed gating procedures for analysis of fetal and postnatal HSPCs. BM mononuclear cells (BMNCs) from a 18-years donor were labeled with a 19-antibody panel. Gates were set on CD45^+^ cells before selection of the CD34^+^Lin^-^ compartment (Lin: CD3, CD14, CD19, CD24, CD56) (upper left density plot). CD34^+^Lin^-^ HSPCs were first fractionated according to differential CD115 and CD123 expression allowing for isolation of CD115^+^CD123^-^ monocyte (Mo) or CD115^-/+^CD123^+^ dendritic cell (DC) precursors. Remaining CD115^-^CD123^-^ cells were further subdivided into CD45RA^-^ or CD45RA^+^ compartments. Within CD45RA^-^ cells, the most immature CD71^-^ITGB7^-^ fraction was subfractionated into CD38^lo/-^ HSCs or CD38^+^ MPPs subsets. The following procedures were used to further subdivide the CD45RA^+^Lin^-^ HPC compartment: <u>. Gating Strategy-1</u> - (*used for dynamic follow-up of age-dependent variations in ELP production patterns and CD7/CD10 expression*): CD45RA^+^ HPCs are fractionated based on CD117 and CD33 expression to distinguish CD117^hi^CD33^lo^ or CD117^int^CD33^lo^ fractions respectively enriched in granulocyte or dendritic precursors, from the CD117^lo/-^CD33^-^ lymphoid progenitors (LyPs). The lymphoid compartment is further subdivided into ITGB7^-^CD127^-^ MLP or CD127^-/+^ ELP subfractions. <u>. Gating Strategy-2</u> - (*used for dynamic follow-up of CD10^+^ MLPs and CD127^+^ ELPs in the BM of postnatal donors*). Lymphoid progenitors (LyP) are subdivided into CD10^+^CD127^-^ MLP or CD127^+^ CD10^-/+^ ELPs fractions. <u>. Gating Strategy-3</u> - (*used for bulk or clonal in vitro diversification assays*). Procedure used for isolation of BM CD45RA^-^CD38^lo^ HSCs (left panel) or CD45RA^+^CD7^-^CD10^-^ LMDPs.

**Extended Data Figure 8.**
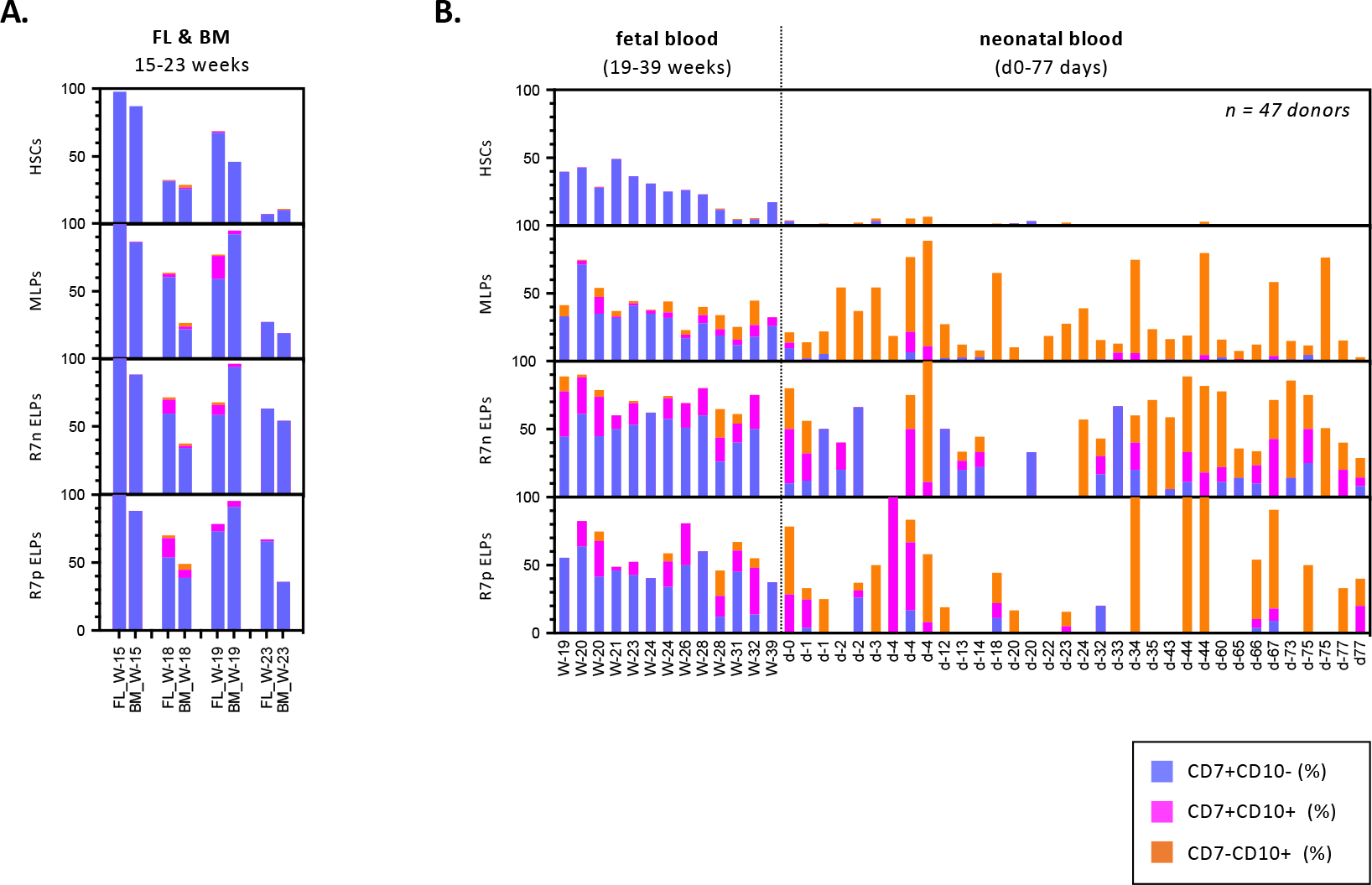
CD7 and CD10 expression patterns by fetal and neonatal HSPCs. (A). Barplots show CD7 and CD10 expression by fetal liver (FL) or BM CD34^+^ HSPCs from 4 donors between 15 and 23 development weeks; gates were set as described above (Extended Data Fig. 7: Gating Strategy-1); results are expressed as relative percentages of CD7 or CD10-expressing cells within the indicated cellular subsets. (B). Barplots show age-dependent variations in CD7 and CD10 expression by CD34^+^ HSPCs circulating in the blood of 47 donors between 19 PCW and 77 days postnatal.

**Extended Data Figure 9.**
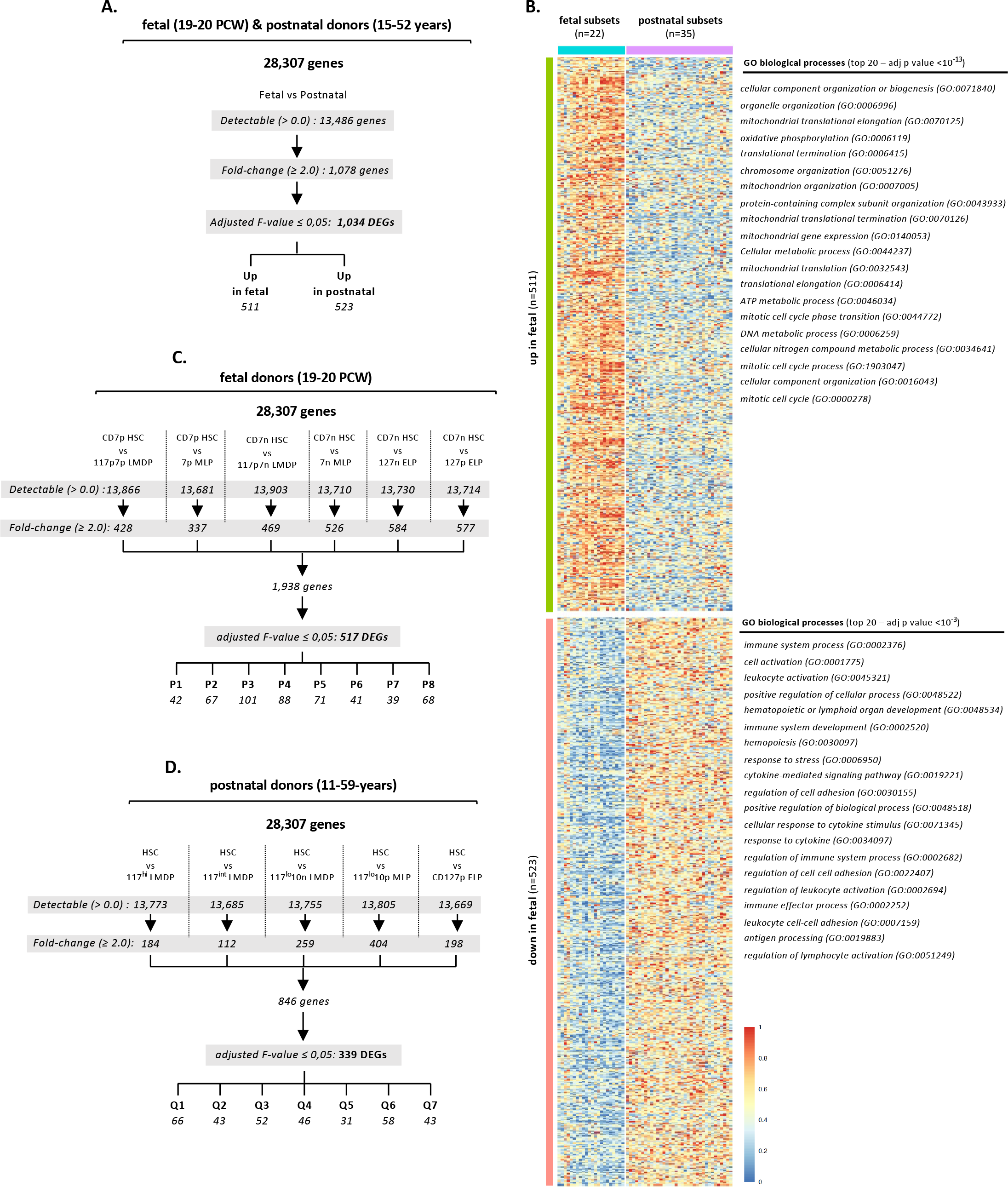
Transcriptional profiling of 8 fetal and 6 postnatal CD34^+^ HSPC subsets. (A). Pipeline based on comparison between fetal and postnatal HSPCs. Amongst 1034 DEGs, 511 were upregulated in fetal HSPCs versus 523 in their postnatal counterparts. (B). Heatmap show clustering and annotations of 1034 DEGs (see also Table S3). Gene Ontology annotations correspond to the top 20 Biological Processes differentially expressed fetal and postnatal HSPCs; adjusted p values are indicated. (C, D). Pipelines based on multiple pairwise comparisons used for extraction of population-specific gene signatures across (C) 8 fetal or (D) 6 postnatal BM HSPC subsets. Detection thresholds, fold-changes and adjusted F-value are indicated (see also Tables S4-5).

**Extended Data Figure 10.**
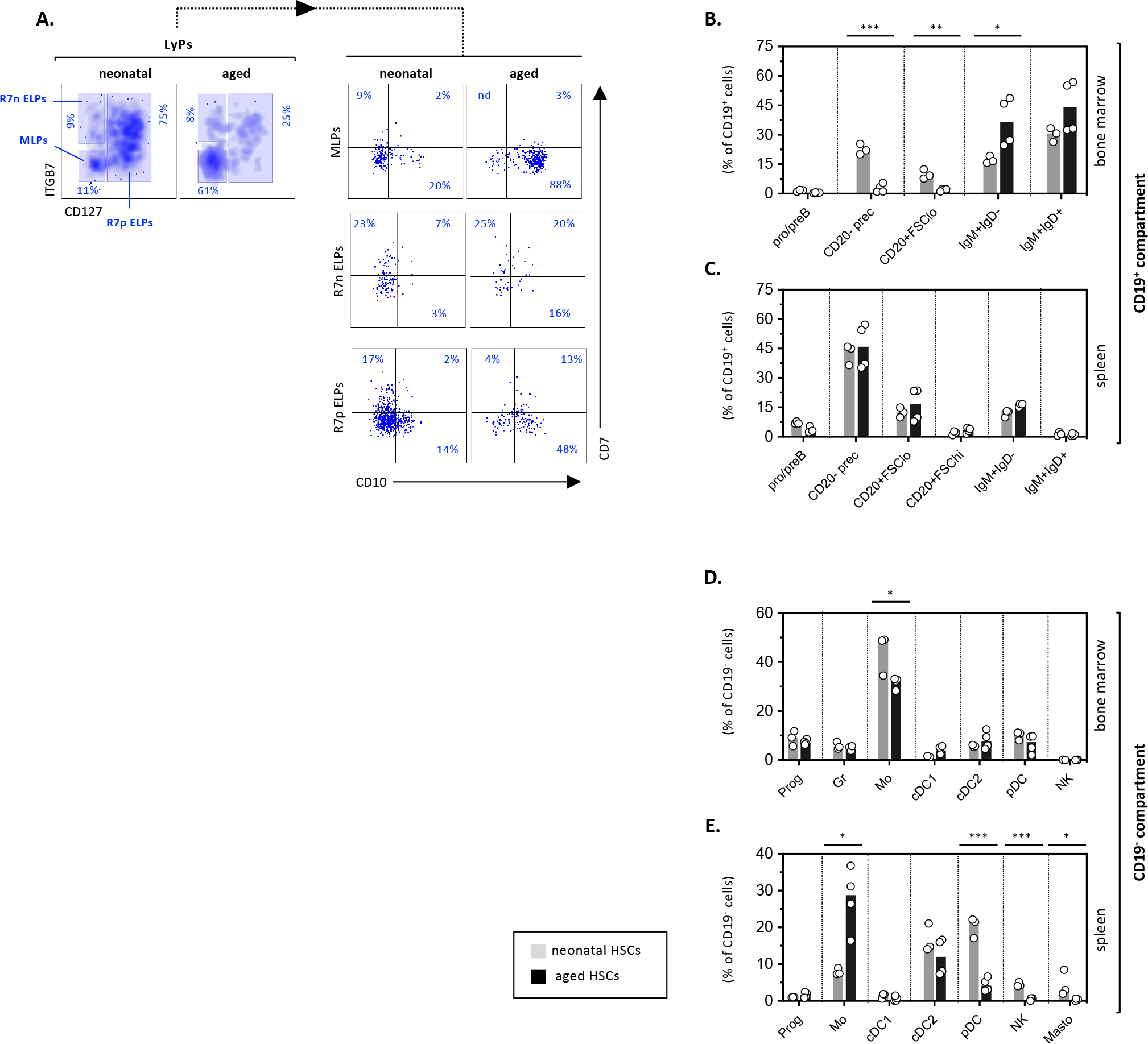
In vivo assessment of the differentiation potential of CD34^+^ HSCs from elderly donors (≥60-years) (A). Immunophenotypic profiling of hu-CD45^+^ BM HSPCs. Mice were sacrificed 8 weeks after intravenous injection of neonatal (nnHSCs: UCB) or aged (aHSCs: 65 or 80 year-old donors) HSCs. BMNCs were labeled with a 32-antibody panel before analysis by spectral cytometry. FACS data are pooled from 3 (nnHSCs) or 4 (aHSCs) xenografted mice. Gates are set on the CD34^+^Lin^-^CD45RA^+^CD117^lo^ Ly compartment. Left density plots show the subdivision of lymphoid progenitors into MLP and CD127^-^ or CD127^+^ ELP subsets; right dot plots show CD7 and CD10 expression; percentages of the corresponding subsets are indicated. (B-E). Immunophenotypic profiling of mature CD19^+^ or CD19^-^ cellular subsets. Hu-CD45^+^ cells isolated as above from the BM or spleen of mice xenografted with neonatal or aged HSCs were analyzed as above. Gates are set on the CD34^-^ compartment followed by subdidivision into CD19^+^ or CD19^-^ compartments. Upper bar plots show stratification of (B) BM or (C) spleen CD19^+^ BLs based on forward scatter characteristic and differential CD20, IgM or IgD expression. Lower bar plots show the relative percentages of the indicated CD56^+^ NK, CD14^+^ monocyte, CD141^+^ cCD1, CD1c^+^ cDC2 or CD117^hi^ mastocyte subsets in (D) BM or (E) spleen; results are normalized relative to total CD19^-^ cells. Circles correspond to individual mice; bars indicate medians; assessment of the statistical significance is based on the unpaired t-test (* p<0.05; *** p<0.001).

**Extended Data Figure 11.**
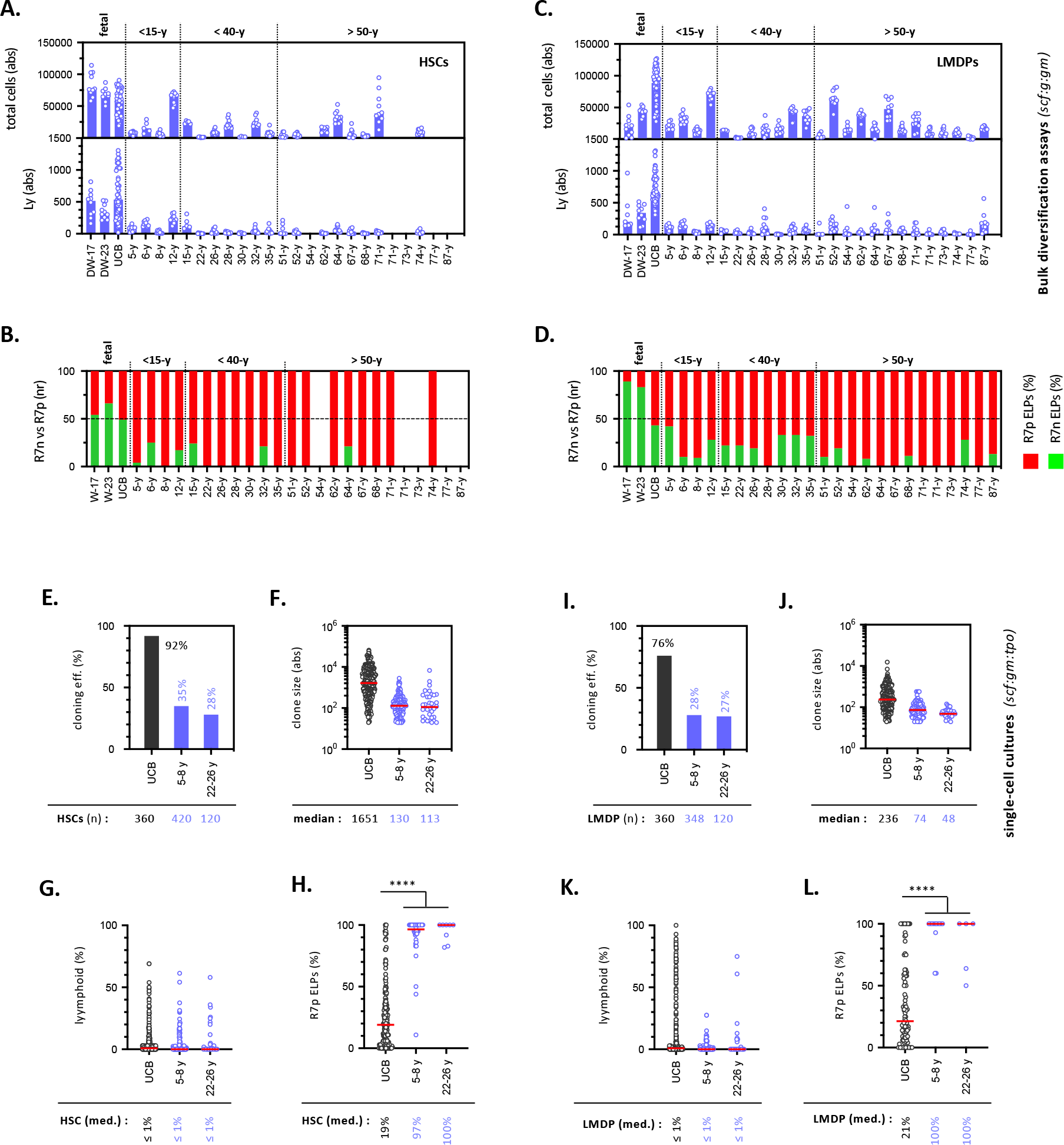
Age-dependent variations in the lymphoid potential of HSCs and LMDPs. (A-D). Assessement of lymphoid potential in bulk diversification assays. HSCs or LMDPs sorted as described in Figure S7 (Gating Strategy-2) from the UCB or the BM of 2 fetal and 24 postnatal donors between development week-17 and 87 years were cultured for 10 days by 100-cell pools under the *scf:g:gm* condition before FACS analysis. <u>. Quantification of HSC lymphoid potential</u>: (A) barplots show absolute cells numbers (upper panel) or percentages of lymphoid cells (lower panel) defined as the sum of the percentages of CD127^-/+^ ELP and CD19^+^ B cells; (B) stacked bar plot show the relative proportions of CD127^-^ (green bars) and CD127^+^ (red bars) ELPs from the same donors; bars correspond to median percentages of ≥10 individual wells/donor. . <u>Quantification of LMDP lymphoid potential</u>: (C) barplots show absolute cell number or percentages of lymphoid cells; and (D) stacked plots show the relative proportion of CD127^-^ and CD127^+^ ELPs. (E-L). Assessement of lymphoid potential in clonal diversification assays. HSCs or LMDPs isolated from the UCB or the BM of 4 postnatal donors between 5-8 years or 22-26 years. Cells were seeded individually in 96-well plates and cultured for 14 days onto OP9 stroma under the *scf:gm:tpo* before FACS analysis. <u>. Quantification of clonal HSC lymphoid output</u>: Bar and circle plots respectively show (E) cloning efficiencies; (F) absolute cell numbers per clone; (G) total lymphoid cell percentage per clone; (H) percentages of CD127^+^ ELPs per clone normalized relative to total ELPs; positivity threshold for ELP detection is set arbitrarily at ≥ 10 ELPs/clone. Red bars correspond to medians; corresponding values are indicated below each plot. Assessment of statistical significance was performed with the Mann-Whitney test (**** p<0.0001; *** p<0.001; ** p<0.01;). <u>. Quantification of clonal LMDP lymphoid output</u> : Bar and circle plots show (I) cloning efficiencies; (J) absolute cell numbers; (K) total lymphoid cell percentages; (L) percentages of CD127^+^ ELPs per clone.

